# A model for physiological transmembrane transport derived from thermodynamical principles

**DOI:** 10.1101/403238

**Authors:** Marco Arieli Herrera-Valdez

## Abstract

A generic formulation for both passive and active transmembrane transport is derived from basic thermodynamical principles. The derivation takes into account the energy required for the motion of molecules across membranes, and includes the possibility of modeling asymmetric flow. Transmembrane currents can then be described by the generic model in the case of electrogenic flow. As it is desirable in new models, it is possible to derive other well known expressions for transmembrane currents as particular cases of the generic formulation. For instance, the conductance-based formulation for current turns out to be a linear approximation of the generic current. Also, under suitable assumptions, other formulas for current based on electrodiffusion, like the constant field approximation by Goldman, can also be recovered from the generic formulation. The applicability of the generic formulations is illustrated first with fits to existing data, and after, with models of transmembrane potential dynamics for pacemaking cardiocytes and neurons. The generic formulations presented here provide a common ground for the biophysical study of physiological phenomena that depend on transmembrane transport.

## Introduction

One of the most important physiological mechanisms underlying communication within and between cells is the transport of molecules across membranes. Molecules can cross membranes either passively (Stein and Litman, 2014), or via active transport (Bennett, 1956). Molecules are passively transported across a membrane when they move along their (electro)chemical gradient and occurs through channels that may be spontaneously formed within the lipid bilayer (Blicher and Heimburg, 2013), or lined by transmembrane proteins (Hille, 1992; Stein and Litman, 2014) that may be selective for molecules of specific types (Favre et al., 1996; Almers and McCleskey, 1984; Doyle et al., 1998). Therefore, passive transport is (electro)diffusive in nature. In contrast, active transport takes molecules against their electrochemical gradients, and is mediated by transmembrane proteins commonly called pumps (e.g. symporters, exchangers) that mechanically translocate the molecules they transport (Bennett, 1956; Ussing, 1949a,b). The energy for active transport of molecules may be obtained from biochemical reactions (e.g. ATPases, light-driven pumps) or from the electrochemical gradients of molecules transported in parallel to the molecule that is actively transported (SKou, 1965). One important functional distinction between channels and pumps is that the rate of transport for channels is generally several orders of magnitude faster than the rate for pump-mediated transport (Ussing, 1949c; Gadsby, 2009). Such differences are reflected in the sizes of different transmembrane currents typically observed in excitable cells (Herrera-Valdez and Lega, 2010).

Theoretical models of transmembrane transport play a critical role in developing our understanding of the function and mechanisms underlying electrical signaling and cellular excitability (Goldman, 1943; Barr, 1965; Cole, 1965; Kell, 1979; Läuger, 1973; Stevens and Tsien, 1979; Wiggins, 1985a,b,c; DiFrancesco and Noble, 1985; Endresen et al., 2000; Gadsby, 2009), and some of its associated pathologies (Marbán, 2002; Ashcroft, 2005). The best known transmembrane transport models include the widely used conductance-based formulation from the seminal work of Hodgkin and Huxley (1952), the Goldman-Hodgkin-Katz equation (Hodgkin and Katz, 1949; Goldman, 1943; Pickard, 1976), and several other expressions for carrier and channel mediated transport with many different functional forms (Rosenberg and Wilbrandt, 1955; Rasmusson et al., 1990a,b; DiFrancesco and Noble, 1985). Other formulations for *ionic* transport across membranes derived from biophysical principles available in the literature include those in the seminal work by Pickard (1969, 1976), Jacquez and Schultz (1974); see also Jacquez (1981) and similar work by Endresen et al. (2000), and those in the excellent book by Johnston et al. (1995). Such formulations describe the relationship between the activity and permeability of ions across membranes, and the transmembrane potential. However general models that describe physiological transport that include passive and active transport of charged or non-charged molecules, possibly including bidirectional but asymmetric flows, are still missing. The work presented here builds upon the results previously mentioned by describing transport macroscopically in terms of the energy required to move molecules across a membrane. The result is a generic formulation with a common functional form for both passive and active transport (Herrera-Valdez, 2014) that also includes a term that regulates the asymmetry in the flow (rectification).

The details of the derivation can be found are explained in the next section. Examples of fits to experimental data and features like asymmetric bidirectional flow. An application of the generic formulation is illustrated with models for the transmembrane potential dynamics in cardiac pacemaker cells and striatal fast spiking interneurons (Appendix) using the same functional forms for the currents. Derivations of formulas and specific examples of noncentral issues addressed in this article can be found in the Appendix.

## Generic formulation for transmembrane flux and current

### Work required for transmembrane molecular fluxes

Consider a system consisting of a biological membrane surrounded by two aqueous compartments (e.g. extracellular and intracellular). Assume, to start with, that the compartments contain molecules of a single type *s* (*e*.*g*. Na^+^, K^+^, glucose), possibly in different concentrations. Let Δ*G*_*s*_ be the energy required for the transport of the molecules across the membrane in a specific direction (e.g. inside to outside). To write an expression for Δ*G*_*s*_ it is necessary to take the direction of motion of the *s*-molecules into account. To do so, label the extracellular and intracellular compartments as 0 and 1, respectively, and let *c*_*s*_ and *d*_*s*_ represent the source and the destination compartments for the transport of the *s*-molecules. The pair (*c*_*s*_, *d*_*s*_)=(0,1) represents inward transport and the pair (*c*_*s*_, *d*_*s*_)=(1,0) represents outward transport. The work required to transport *n*_*s*_ molecules of type *s* from compartment *c*_*s*_ to compartment *d*_*s*_ can then be written as

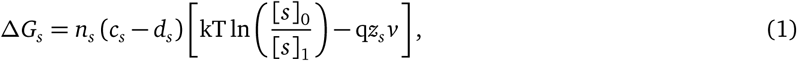

(Weer et al., 1988; Aidley, 1998; Blaustein et al., 2004) where q, *z*_*s*_, [*s*]_0_, and [*s*]_1_ represent the elementary charge, the valence, the extracellular, and the intracellular concentrations for the molecules of type *s*, respectively. Two particular cases are worth noticing. First, if *s* is an ion, then *z*_*s*_ *≠* 0 and equation (1) becomes

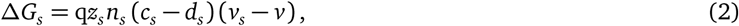

where *v*_*s*_ is the Nernst potential for the *s*-molecules^1^ (Nernst, 1888). Second, if the *s*-molecules are not charged, then *z*_*s*_ = 0 and the work required to move the *s*-type molecules from *c*_*s*_ to *d*_*s*_ simplifies to

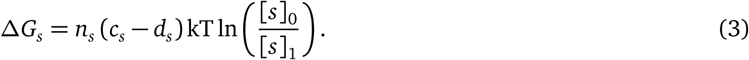

If Δ*G*_*s*_ < 0, then the molecules can be transported passively (*e*.*g*. electrodiffusion), decreasing the electrochemical gradient for *s* across the membrane. In contrast, if Δ*G*_*s*_ > 0, the transmembrane transport of *s* from *c*_*s*_ to *d*_*s*_ is not thermodynamically favorable, which means the transport from *c*_*s*_ to *d*_*s*_ requires energy that is not available in the electrochemical gradient for *s* (active transport). As a consequence, active transport of *s* would increase the driving force for the motion of *s* across the membrane.

### Joint transmembrane transport of different types of molecules

To find an expression for Δ*G* that describes a more general transport mechanism, assume that transport takes place as single events in which molecules of *m* different types move in parallel, or possibly sequentially (e.g. first Na^+^, then K^+^), across the membrane. Let *S* be a set that represents the types of molecules that are jointly transported in a single event. For instance, for Na^+^-H^+^ exchangers, *S* = {*Na*, *H*}, with *m* = 2. The energy required to transport these molecules is the sum of the energies required to transport each of the molecules in *S*. In other words,

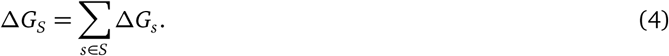

As before, transport is thermodynamically favorable when Δ*G*_*S*_ ≤ 0. If not, extra energy is required. To distinguish between these two cases, define the total energy of the transport mechanism as

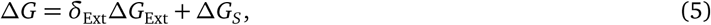

where δ_Ext_ = 1 if Δ*G*_*S*_ > 0, and 0 otherwise. In particular, for ATP-driven transport, the extra energy supplied by hydrolysis of ATP (Tanford, 1981; Weer et al., 1988) is

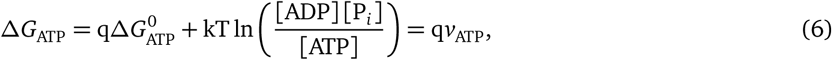

where *v*_ATP_ *≈-* 450 mV (Endresen et al., 2000), but could vary depending on the amounts of ATP, ADP, and P_*i*_ (Weer et al., 1988). *Similar expressions could be derived for active transport driven by light*, *or other sources of energy*.

### Flux due to transmembrane transport

The formulation in equation (10) can be combined with equation (1) to derive a generalized expression for flux and model different known mechanisms of physiological transmembrane transport, possibly combining the transport of different molecules simultaneously (e.g. Na-H exchange). In this case, the forward direction of the transport would be described by the combined forward transport of each of the different molecules under consideration. For instance, the source and target compartments for Na^*+*^ and *Ca*^*2+*^ are different in Na-Ca exchangers. The stoichiometry for the transport mediated by Na-Ca exchangers in the forward direction involves three Na^*+*^ molecules moving inward (along their electrochemical gradient) in exchange for one Ca^*2+*^ molecule moving outward (against their electrochemical gradient) (Mullins, 1979; Venetucci et al., 2007).

*Let α* and *β* be the flux rates in the forward and backward directions, in units of molecules per ms per *µ*m^*-*2^. These rates depend, a priori, on the energy required for the transport of the molecules in *S*. The net *flux* rate associated to the net transmembrane transport, can then be written as

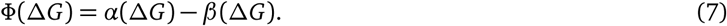

How do *α* and *β* depend on Δ*G*? The steady state relationship between the energy Δ*G* and the the forward and backward flow rates, hereby represented by *α* and *β*, can be written as

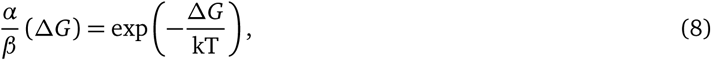

where k is Boltzmann’s constant, and T the absolute temperature.

Assuming that *α* and *β* are continuous functions, the rates *α* and *β* can be rewritten as

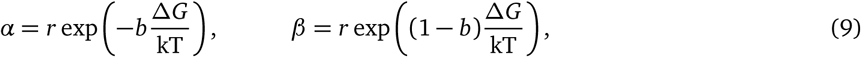

where *r* (molecules per ms per *µ*m^*-*2^) may depend on temperature (Sen and Widdas, 1962), the transmembrane potential (Starace et al., 1997), the concentrations inside and outside of the membrane (Yue et al., 1990), and other factors (Novák and Tyson, 2008). Note that the functional form of the rates in equations (9) are similar to those by Butler (1924); Erdey-Grúz and Volmer (1930). Also, notice that the steady state relationship between *α* and *β* in equation (8) can be obtained from equations (9), for any *r* and any *b*. However, it should be the case that *r* and *b* vary in specific ranges depending on the physico-chemical characteristics of the pore through which molecules cross the membrane, and in general, on the the transport mechanism. As mentioned earlier, the rate *r* should larger for electrodiffusive transport in comparison to the slower transport rates for pumps and other carrier proteins. If the parameter *b ∈* [0, 1], then *b*Δ*G* and (*b-* 1)Δ*G* have opposite signs and can be thought of as the energies required to the transport of the molecules in *S* in the forward and backward directions, respectively, with *b* biasing the transport in the forward direction when close to 1, and in the backward direction when close to 0 (Fig. 1).

**Figure 1.**
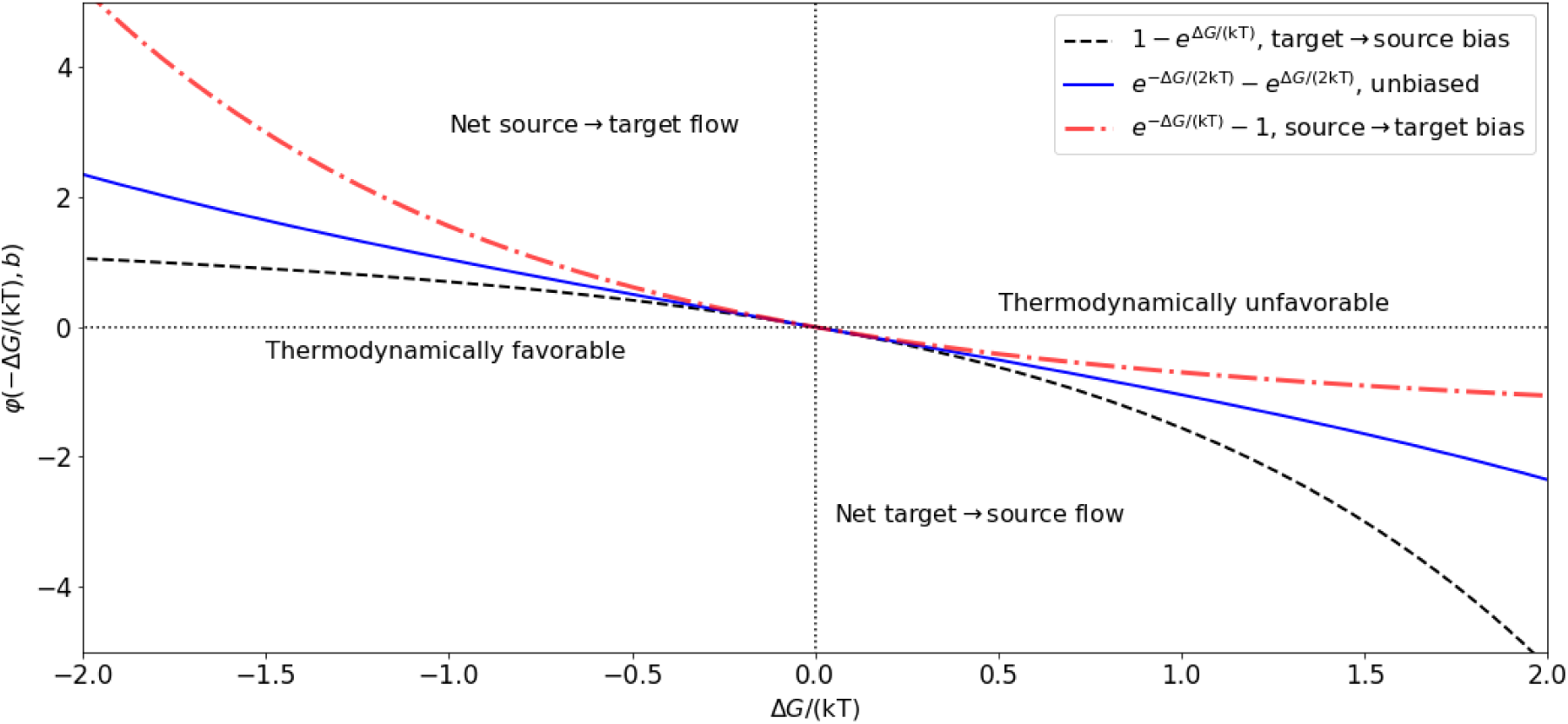
**Fluxes biased in the target** → **source (backward,** *b***=0.1, black dashed line), source** → **target (forward,** *b* **=0.9, red dash-dot line), or showing no rectification (***b***=0.5, blue solid line**). See Table 1 for examples.

**Table 1.** Energy required for transmembrane transport mediated by different passive and active mechanisms.

| Pump or channel | Molecule (s) | $n_s$ | $c_s$ | $d_s$ | $c_s - d_s$ | $\Delta G_s = qn_s(c_s - d_s) \left[ kT \log \left( \frac{[s]_0}{[s]_1} \right) - qz_s v \right]$ | $\eta$ | $v_o$ | $\alpha/\beta = \exp \left( -\frac{\Delta G}{kT} \right)$ |
| --- | --- | --- | --- | --- | --- | --- | --- | --- | --- |
| $\text{Cl}^-$ channel | $\text{Cl}^-$ | 1 | 0 | 1 | -1 | $\Delta G_{\text{Cl}} = q(v_{\text{Cl}} - v)$ | 1 | $v_{\text{Cl}}$ | $\left( \frac{[\text{Cl}]_1}{[\text{Cl}]_0} \right) \exp \left( \frac{v}{v_T} \right)$ |
| $\text{K}^+$ channel | $\text{K}^+$ | 1 | 1 | 0 | 1 | $\Delta G_{\text{K}} = q(v_{\text{K}} - v)$ | 1 | $v_{\text{K}}$ | $\left( \frac{[\text{K}]_1}{[\text{K}]_0} \right) \exp \left( \frac{v}{v_T} \right)$ |
| $\text{Na}^+$ channel | $\text{Na}^+$ | 1 | 0 | 1 | -1 | $\Delta G_{\text{Na}} = -q(v_{\text{Na}} - v)$ | -1 | $-v_{\text{Na}}$ | $\left( \frac{[\text{Na}]_0}{[\text{Na}]_1} \right) \exp \left( -\frac{v}{v_T} \right)$ |
| $\text{Ca}^{2+}$ channel | $\text{Ca}^{2+}$ | 1 | 0 | 1 | -1 | $\Delta G_{\text{Ca}} = -2q(v_{\text{Ca}} - v)$ | -2 | $-2v_{\text{Ca}}$ | $\left( \frac{[\text{Ca}]_0}{[\text{Ca}]_1} \right) \exp \left( -2\frac{v}{v_T} \right)$ |
| $\text{Na}^+ - \text{K}^+$ ATPase | $\text{Na}^+$<br>$\text{K}^+$ | 3<br>2 | 1<br>0 | 0<br>1 | 1<br>-1 | $\Delta G_{\text{Na}} = 3q(v_{\text{Na}} - v)$<br>$\Delta G_{\text{K}} = -2q(v_{\text{K}} - v)$ | 1 | $v_{\text{ATP}} + 3v_{\text{Na}} - 2v_{\text{K}}$ | $\left( \frac{[\text{Na}]_1}{[\text{Na}]_0} \right)^3 \left( \frac{[\text{K}]_0}{[\text{K}]_1} \right)^2 \exp \left( \frac{v - v_{\text{ATP}}}{v_T} \right)$ |
| $\text{Ca}^{2+}$ ATPase | $\text{Ca}^{2+}$ | 1 | 1 | 0 | 1 | $\Delta G_{\text{Ca}} = 2q(v_{\text{Ca}} - v)$ | 2 | $v_{\text{ATP}} + 2v_{\text{Ca}}$ | $\left( \frac{[\text{Ca}]_1}{[\text{Ca}]_0} \right) \exp \left( \frac{2v - v_{\text{ATP}}}{v_T} \right)$ |
| $\text{H}^+$ ATPase | $\text{H}^+$ | 1 | 1 | 0 | 1 | $\Delta G_{\text{H}} = q(v_{\text{H}} - v)$ | 1 | $v_{\text{ATP}} + v_{\text{H}}$ | $\left( \frac{[\text{H}]_1}{[\text{H}]_0} \right) \exp \left( \frac{v - v_{\text{ATP}}}{v_T} \right)$ |
| $\text{Na}^+ - \text{Ca}^{2+}$ exchanger | $\text{Na}^+$<br>$\text{Ca}^{2+}$ | 3<br>1 | 0<br>1 | 1<br>0 | -1<br>1 | $\Delta G_{\text{Na}} = -3q(v_{\text{Na}} - v)$<br>$\Delta G_{\text{Ca}} = 2q(v_{\text{Ca}} - v)$ | -1 | $2v_{\text{Ca}} - 3v_{\text{Na}}$ | $\left( \frac{[\text{Na}]_1}{[\text{Na}]_0} \right)^3 \left( \frac{[\text{Ca}]_0}{[\text{Ca}]_1} \right) \exp \left( -\frac{v}{v_T} \right)$ |
| $\text{Na}^+ - \text{I}^-$ symporter | $\text{Na}^+$<br>$\text{I}^-$ | 2<br>1 | 0<br>0 | 1<br>1 | -1<br>-1 | $\Delta G_{\text{Na}} = -2q(v_{\text{Na}} - v)$<br>$\Delta G_{\text{I}} = -q(v_{\text{I}} - v)$ | -1 | $-v_{\text{I}} - 2v_{\text{Na}}$ | $\left( \frac{[\text{Na}]_1}{[\text{Na}]_0} \right)^2 \left( \frac{[\text{I}]_1}{[\text{I}]_0} \right) \exp \left( -\frac{v}{v_T} \right)$ |
| $\text{Na}^+ - \text{H}^+$ exchanger | $\text{Na}^+$<br>$\text{H}^+$ | 1<br>1 | 0<br>1 | 1<br>0 | -1<br>1 | $\Delta G_{\text{Na}} = -q(v_{\text{Na}} - v)$<br>$\Delta G_{\text{H}} = q(v_{\text{H}} - v)$ | 0 | $v_{\text{H}} - v_{\text{Na}}$ | $\left( \frac{[\text{H}]_1}{[\text{H}]_0} \right) \left( \frac{[\text{Na}]_0}{[\text{Na}]_1} \right)$ |
| $\text{K}^+ - \text{Cl}^-$ symporter | $\text{K}^+$<br>$\text{Cl}^-$ | 1<br>1 | 1<br>1 | 0<br>0 | 1<br>1 | $\Delta G_{\text{K}} = q(v_{\text{K}} - v)$<br>$\Delta G_{\text{Cl}} = -q(v_{\text{Cl}} - v)$ | 0 | $v_{\text{K}} - v_{\text{Cl}}$ | $\left( \frac{[\text{K}]_1}{[\text{K}]_0} \right) \left( \frac{[\text{Cl}]_1}{[\text{Cl}]_0} \right)$ |
| $\text{Na}^+ - \text{K}^+ - \text{Cl}^-$ symporter | $\text{Na}^+$<br>$\text{K}^+$<br>$\text{Cl}^-$ | 1<br>1<br>2 | 0<br>0<br>0 | 1<br>1<br>1 | -1<br>-1<br>-1 | $\Delta G_{\text{Na}} = -q(v_{\text{Na}} - v)$<br>$\Delta G_{\text{K}} = -q(v_{\text{K}} - v)$<br>$\Delta G_{\text{Cl}} = 2q(v_{\text{Cl}} - v)$ | 0 | $2v_{\text{Cl}} - v_{\text{Na}} - v_{\text{K}}$ | $\left( \frac{[\text{Na}]_0}{[\text{Na}]_1} \right) \left( \frac{[\text{K}]_0}{[\text{K}]_1} \right) \left( \frac{[\text{Cl}]_1}{[\text{Cl}]_0} \right)^2$ |

The flux can then be written explicitly combining equations (7) and (9) to obtain,

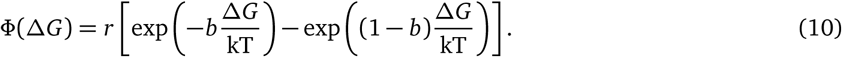

Taking the above observations into account, it is possible to combine equations (4) and (5), to write an expression similar to equation (8) for the steady state balance between the forward and backward transport of all the molecules in *S*.

### Flux and current

Substitution of the formulas for Δ*G* from equations (4) (5) into equation (10), the flux rate resulting from simultaneously transporting molecules in *S* across the membrane can be written explicitly

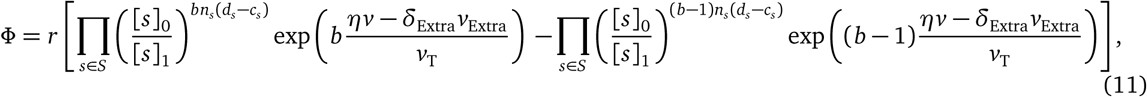

where *v*_T_ = kT/q and

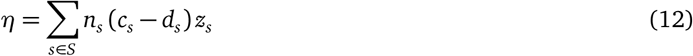

represents the net number of charges moved across the membrane.

If the transport is electrogenic, then the product q*η* (in Coulombs) represents the net charge moved across the membrane, relative to the extracellular compartment. Non electrogenic transport yields *η* = 0, which means the flow does not depend on the transmembrane potential, and

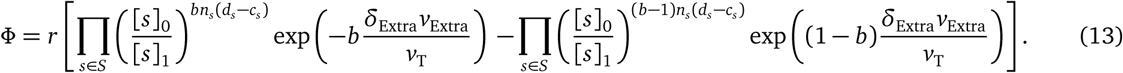

If only ions are involved in the transport, the flux simplifies to

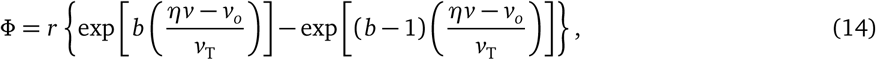

where

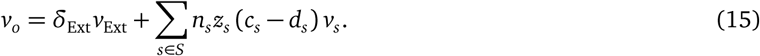

The quantity *v*_*o*_*/η* can be thought of as a reversal potential. If *η <* 0, then positive charge is transported inward, or negative charge is transported outward. In contrast, *η >* 0 means that positive charge is transported outward or negative charge transported inward. For instance, inward electrodiffusion of single Na^+^ ions gives an *η* = −1, which can be thought of as loosing one positive charge from the extracellular compartment in each transport event (see Table 1). In particular, for electrodiffusive (passive) transport of ions of type *l*, *v*_*o*_ reduces to *n*_*l*_ *z*_*l*_ (*c*_*l*_ − *d*_*l*_) *v*_*l*_. A list with examples of energies and total charge movements for different transport mechanisms can be found Table 1.

The first, more complex, form of the flux in equation (13) could be useful when working with models for which changes in the concentrations of different molecules are relevant.

#### Transmembrane current

The flux that results in electrogenic transport (equations (13) and (14)) can be converted to current density after multiplication by q*η*. In short form,

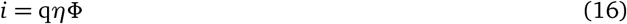

with q*r* in amperes/m^2^ or equivalent units.

Substitution of equations (13) or (14) into equation (16) yields a general formula for the current generated by transmembrane ionic flux (Fig. 2), that uses the same functional form for channels (protein or lipid) and pumps. Recall that equation (16) can also be written explicitly in terms of the transmembrane concentrations of one or more of the ions involved using equation (13). It is possible to derive expressions for *r* that take into account biophysical variables like temperature and the shape and length of the pore through which the molecules cross (Pickard, 1969; Endresen et al., 2000).

**Figure 2.**
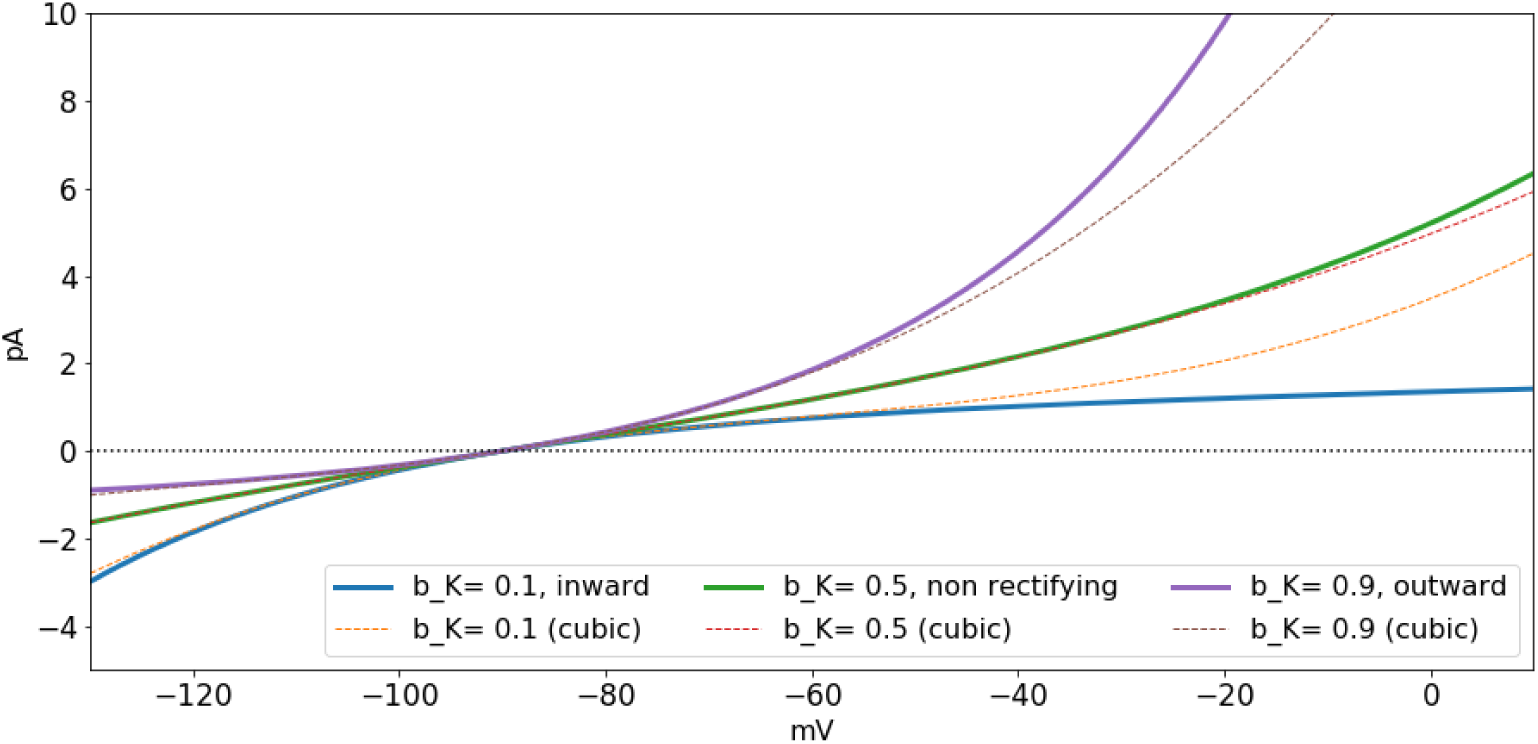
**Fluxes for K-electrodiffusion for** *b*_K_ *∈*{0.1, 0.5, 0.9}**and their cubic approximations. Inward rectification occurs for** *b*_*K*_ < 1/2 **and outward rectification for** *b*_K_ > 1/2 **and** q*r*_K_ *N*_K_ = 1.

### Special cases and examples

A number of nontrivial and important properties of transmembrane ionic currents, including rectification, are also described by equation (16). Also, different models for current already in the literature can be obtained by making approximations or setting particular cases from equation (16). Examples include electrodiffusive currents that result from integration of the Nernst-Planck equation along the length of membrane pore (Pickard, 1969; Jacquez and Schultz, 1974; Johnston et al., 1995). Of particular interest, conductance-based currents are linear approximations of the formulation (16), around the reversal potential for the current.

#### Lower order approximations to the generic formulation and conductance based models

Conductancebased currents (Hodgkin and Huxley, 1952) are linear approximations of the generic current from equation (16), around the reversal potential *v*_*o*_*/η*. To see this, use Taylor’s theorem (Courant and John, 2012; Spivak, 2018) to rewrite the generic current from equation (16) as a series around *v*_*o*_

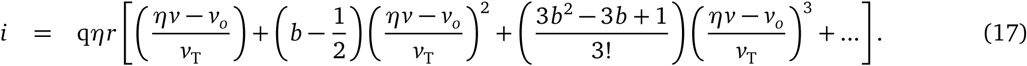

Truncation of the series to first order gives

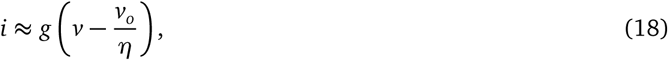

where *g* = *η*^2^q*r/v*_T_ is in units of nS/*µ*m^2^, which has the functional form of the conductance-based current used in the Hodgkin and Huxley (1952) model. For instance, the linear approximation for the current through an open sodium channels around *v*_Na_ in equation (18) gives *g*_Na_ = q*r*_Na_*/v*_T_, and *v*_*o*_ = *η*_Na_ *v*_Na_, with *η*_Na_ = − 1, so that *i*_Na_ *≈g*_Na_(*v-v*_Na_).

Notice that third order approximations to equation (14) can also capture rectification. In contrast, first order approximations (conductance-based models) cannot capture rectification.

#### Rectification results from asymmetric bidirectional flow

The flux of molecules across the membrane can be biased in either the outward or the inward direction when mediated by proteins. This was first called “anomalous rectification” by Katz (1949), who noticed that K^+^ flows through muscle membranes more easily in the inward, than in the outward direction (Armstrong and Binstock, 1965; Adrian, 1969). It was later found some K^+^ channels display the bias in the opposite direction (Woodbury, 1971). The former type of K^+^ current rectification is called inward, and the latter outward.

Rectification is a bias in either of the two directions of transport, which may result from changes in the structure of the proteins or pores through which the molecules cross the membrane (Riedelsberger et al., 2015; Hollmann et al., 1991). The type of rectification (inward or outward) depends on what molecules are being transported and on the structure of the proteins mediating the transport. Rectification is therefore not only a property of ions, as shown by molecules like glucose, which may cross membranes via GLUT transporters bidirectionally, but asymmetrically, even when the glucose concentration is balanced across the membrane (Lowe and Walmsley, 1986).

Rectification can be described by equation (13) by setting *b* to values different from 1/2, and becomes more pronounced as *b* is closer to either 0 or 1. These values represent biases in the transport toward the source, or the target compartment, respectively. As a consequence, rectification yields an asymmetry in the graph of *α - β* as a function of Δ*G* (Fig. 2). For electrogenic transport, rectification can be thought of as an asymmetric relationship between current flow and voltage, with respect to the reversal potential *v*_*o*_. The particular case *b* = 1/2 (non rectifying) yields a functional form for current similar to that proposed by Pickard (1969), and later reproduced by (Endresen et al., 2000), namely

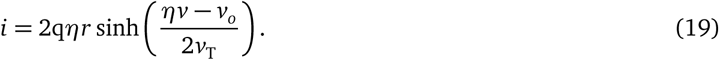

From here on, subscripts will be used to represent different transport mechanisms. For instance, the current for a Na-Ca pump will be written as *i*_NaCa_.

Electrodiffusion of K^+^ through channels (*η* = 1 and *v*_*o*_ = *v*_K_), is outward for *v > v*_K_, and inward for *v < v*_K_. The K^+^ current through the open pore is therefore

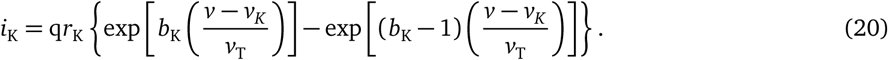

Current flow through inward rectifier channels (Riedelsberger et al., 2015) can be fit to values of *b*_K_ < 1/2. For instance,

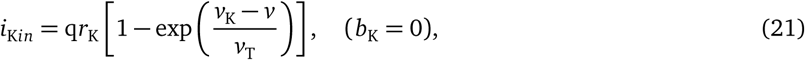

describes a current with limited flow of K^+^ in the outward direction, similar to the currents described originally by Katz (1949). Analogously, *b*_K_ > 1/2 limits the inward flow. For example, the current

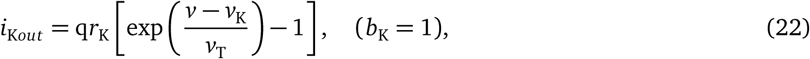

describes outward rectification (Riedelsberger et al., 2015).

Based on the work of Riedelsberger et al. (2015) on K^+^ channels, inward (outward) rectification arises when the S4 segment in K^+^ channels is located in the inner (outer) portion of the membrane. These two generic configurations can be thought of in terms of ranges for the parameter *b*_K_, namely, *b*_K_ < 1/2 for inward, and *b*_K_ > 1/2 for outward rectification.

In general, ion channels are typically formed by different subunits, that may combine in different ways, resulting in structural changes that may restrict the flow of ions through them, causing rectification. For instance, non-NMDA glutamatergic receptors that can be activated by kainic acid and *α*-amino-3-hydroxy-5-methyl-4-isoxazole propionic acid (AMPA) conduct Na^+^, K^+^, and Ca^2+^, with different permeabilities depending on the subunits that form the receptor (Hollmann et al., 1991). The reason is that the specific combination of GluR subunits forming the receptor restrict ionic flow in different ways. In particular, the currents recorded in oocytes injected with combinations of GluR1 and GluR3 cRNA have different steady state amplitudes and show different levels of rectification (Fig. 3).

**Figure 3.**
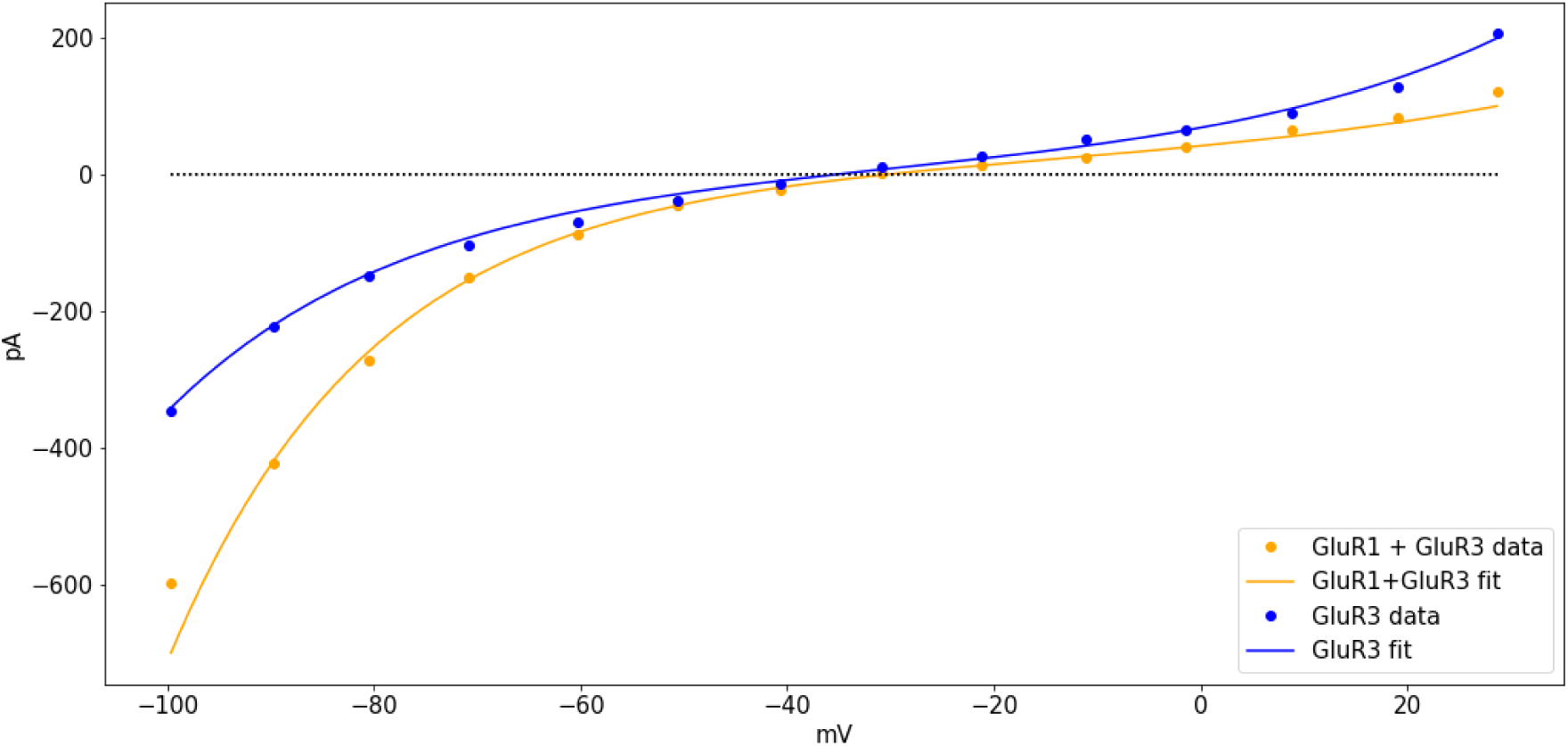
**Currents recorded from oocytes injected with GluR3 cRNA (blue), or a combination of GluR1 and GluR3 cRNA (orange), in Ca**^2+^ **ringer solution after activation by AMPA. The curves were fit with** (*v*_*o*_, *b*, *r*q)**=(- 30,0.45,21), for GluR3 and** (*v*_*o*_, *b*, *r*q)**=(−35,0.35,20), for GluR1+GluR3. The data was digitized from figure 3B in the article by Hollmann et al. (1991), using the *ginput* function from the python module matplotlib (Hunter, 2007)**.

#### Primary active transport

The **Na-K ATPase** is a primary active transporter that uses the energy from the hydrolysis of one molecule of ATP for the uphill transport of Na^+^ and K^+^ (Weer et al., 1988). The kinetics of the **Na-K ATPase** can be assumed to translocate 3 Na ^+^ ions outward and 2 K^+^ ions inward (*η*_NaK_ = 1) with a reversal potential *v*_NaK_ = *v*_ATP_ + 3*v*_Na_ *-*2*v*_K_ (see Table 1) in a single transport event (Post and Jolly, 1957; Garrahan and Glynn, 1967; Gadsby et al., 1985; Chapman, 1973). Importantly, the transport kinetics of the Na-K ATPase and by extension, the current, reverse for potentials smaller than *v*_NaK_ (Weer et al., 1988).

The current-voltage relationships recorded from Na-K ATPases in guinea pig ventricular cells are shaped as hyperbolic sines (Gadsby et al., 1985). Those currents would be fit with *b*_NaK_ *≈*1/2, yielding currents of the form

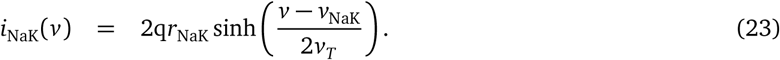

The voltage-dependence of the Na-K ATPase currents is reported to show a plateau as *v* increases past the reversal potential for the current, in response to steroids like strophandin (Nakao and Gadsby, 1989). In such cases, the Na-K ATPase current can be assumed to be inwardly rectifying and fit with values of *b*_NaK_ *≈* 0, so that,

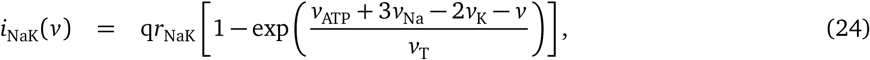

or alternatively,

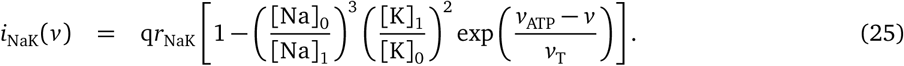

The rectification for the Na-K pump ATPase has also been reported to occur in small neurons of the dorsal root ganglion in rats (Hamada et al., 2003). The alternative expression (25) also explains qualitatively different behaviors of the Na-K current as a function of the transmembrane concentrations of Na^+^ and K^+^. For instance, if either [Na]_1_ or [K]_0_ increase and *v* > *v*_NaK_, then the amplitude of *i*_NaK_ would increase at a smaller rate of change in comparison to when *v* < *v*_NaK_, which grows exponentially in size. This is also in line with reports of non significant changes in the transport by Na-K ATPases in response to elevated intracellular Na^+^ during heart failure (Despa et al., 2002), in which the transmembrane potential is likely to be depolarized.

#### Secondary active transport

An example of a pump that mediates secondary active transport is the *Na-Ca exchanger*, which takes 3 Na^+^ ions from the extracellular compartment in exchange for one intracellular Ca^2+^ ion in forward mode (Pitts, 1979; Reeves and Hale, 1984). The reversal potential for the current is *v*_NaCa_ = 2*v*_Ca_ *-*3*v*_Na_, with *η*_NaCa_ = 1. Assuming *b*_NaCa_ = 1/2, the Na-Ca current is

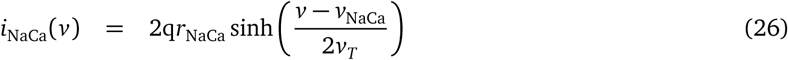

The driving force *v-v*_NaCa_ could reverse in sign with large enough increases in the intracellular concentration of Ca^2+^, or in the membrane potential. As a result, the current could have a dual contribution to the change in transmembrane potential, as predicted by some theoretical models of cardiac pacemaker activity (Rasmusson et al., 1990a,b).

#### Electrodiffusive transport

Consider transmembrane electrodiffusive transport of a single ionic type *x*, with *z*_*x*_ and *v*_*x*_ representing the valence and the Nernst potential for *x*-ions, respectively. In this case, the reversal potential satisfies

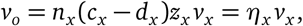

and the generic expression (16) can be rewritten as

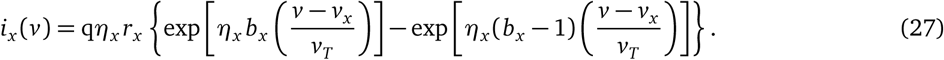

In the absence of rectification (*b*_*x*_ = 0.5),

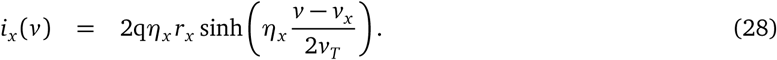

For calcium channels,

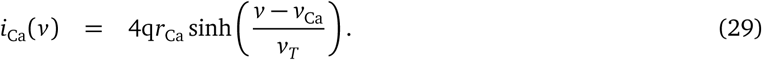

See Pickard (1969); Jacquez and Schultz (1974) Table 1 for other examples.

The applicability of the general formulations described above is illustrated next with models of cardiac and neuronal membrane potential.

## Transmembrane potential dynamics

To show the application of the formulations discussed earlier, let us build a generic model of transmembrane potential dynamics with currents generated by *N* different electrogenic transport mechanisms. For simplification purposes, consider only one such mechanism, labeled as *l*, with *p*_*l*_ *N*_*l*_ active sites, where *N*_*l*_ is the number of membrane sites where the *l*th transport mechanism is found, and *p*_*l*_ is the proportion of active sites (might be voltage or ligand dependent). Then the total current mediated by the *l*th mechanism in a patch of membrane can be written as 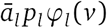 with 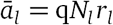 (in pA/μm^2^), and

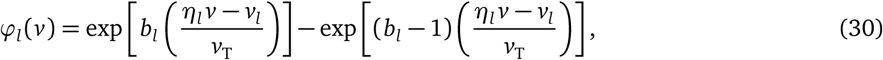

where *v*_*l*_ */η*_*l*_ is the reversal potential for the *l*th current, *l∈* {1, …, *N*} There is experimental evidence for some ion channels that supports the replacement of ā_*l*_ as a constant (Nonner and Eisenberg, 1998). The time-dependent change in transmembrane potential can written as

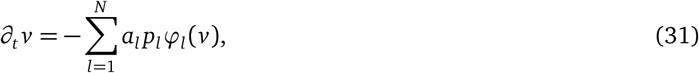

with *v* is in mV and 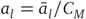 in mV/mS (pA/pF) represents the current amplitude for the *l*th transport mechanism, normalized by the membrane capacitance, for *l*∈ (1, …, *N*). Only electrogenic transport mechanisms are included.

### Cardiac pacemaking in the sinoatrial node

The pacemaking dynamics of cells in the rabbit sinoanatrial node (Fig. 4) can be modeled using low dimensional dynamical systems based on the assumption that *v* changes as a function of K^+^, Ca^2+^, and some Na^+^ transmembrane transport, (Herrera-Valdez and Lega, 2010; Herrera-Valdez, 2014). Transmembrane currents are assumed to be mediated by a combination of channel-mediated electrodiffusion and pumping mechanisms. Explicitly, Ca^2+^ transport is mediated by L-type Ca_*v*13_ channels (Mangoni et al., 2003) and Na^+^-Ca^2+^ exchangers (Sanders et al., 2006). K^+^ transport is mediated by delayed-rectifier voltage-activated channels (Shibasaki, 1987), and Na^+^-K^+^ ATPases (Herrera-Valdez and Lega, 2010; Herrera-Valdez, 2014). In this model, the activation for the L-type Ca^2+^ channels is fast, and assumed to be at steady state (Herrera-Valdez and Lega, 2010). The proportion of activated K^+^ channels and the proportion of inactivated Ca^2+^ channels are both represented by a variable *w* (Herrera-Valdez *and Lega, 2010; Av-Ron et al., 1991). The activation phase of currents recorded in voltage-clamp experiments often displays sigmoidal time courses (Hodgkin and Huxley, 1952; Tsunoda and Salkoff, 1995; Covarrubias et al., 1991). Therefore, the activation dynamics represented by w are described by solutions to equations of the form*

**Figure 4.**
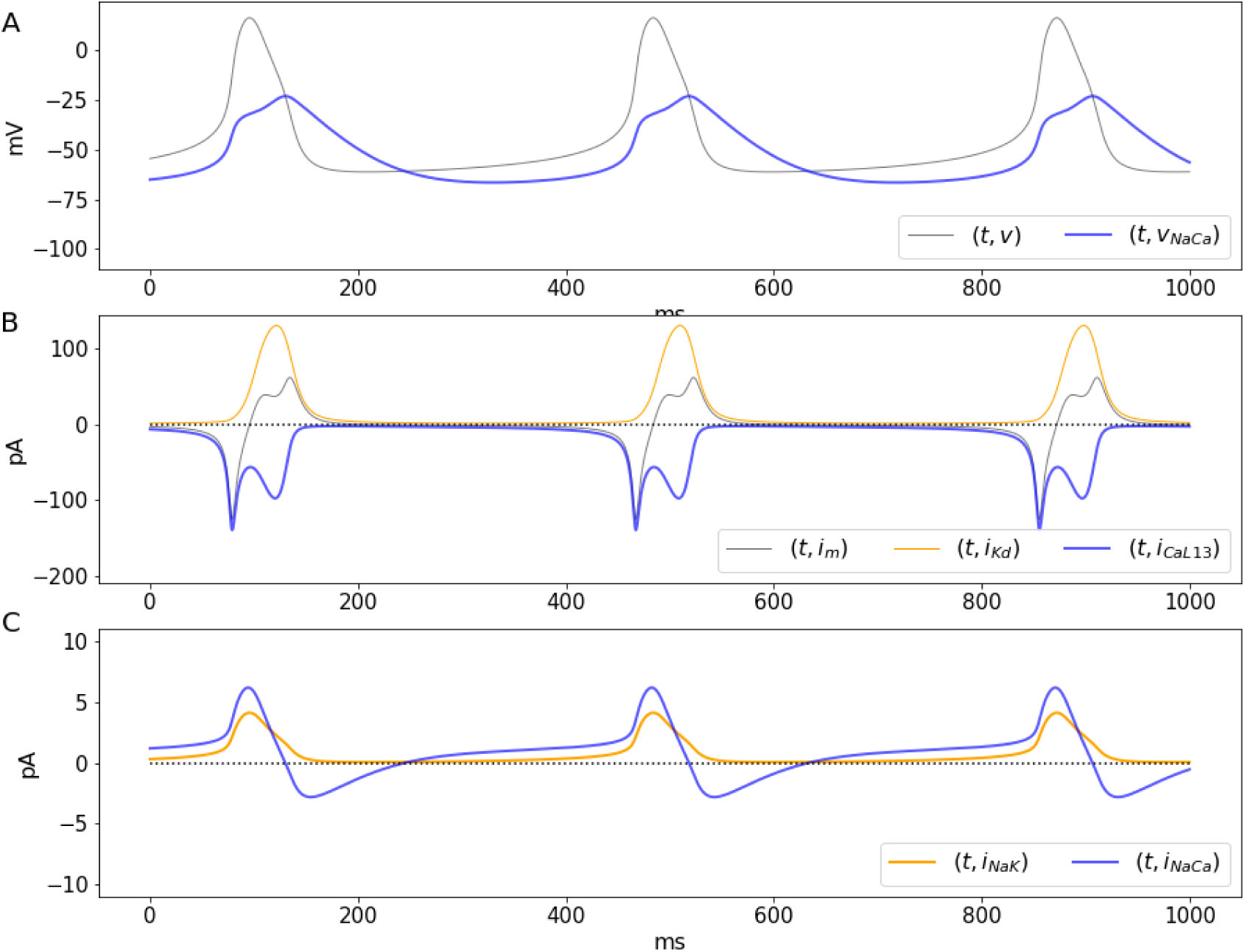
**Central sinoatrial node pacemaking dynamics using the system** (39)**-**(34)**. A. Transmembrane potential and the reversal potential** *v*_NaCa_ **as a function of time. B,C. Dynamics of large currents and small currents, respectively.**

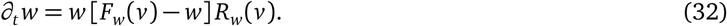

*where F*_*w*_ and *R*_*w*_ represent the voltage-dependent steady state and rate (1/ms) for the opening of Kd channels (Willms et al., 1999). The steady state for the activation of voltage-dependent channels is described by the function

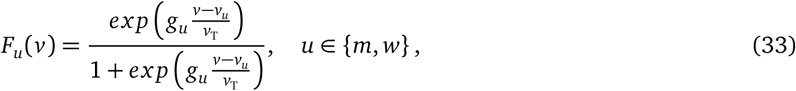

which has a graph with increasing sigmoidal shape as a function of *v*. The parameters *g*_*u*_ and *v*_*u*_ control the steepness and the half-activation potential, for *u* ∈ {*m*, *w*}. The activation rate for K^+^ channels is a voltagen dependent function of the form

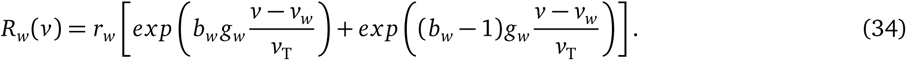

where *b*_*w*_ represents a bias in the conformational change for activation. The function *R*_*w*_ has the shape of a hyperbolic cosine when *b*_*w*_ is 1/2. The transmembrane (normalized) currents are given by

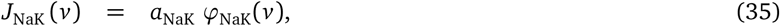

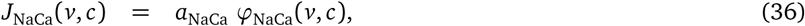

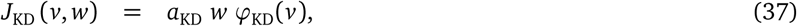

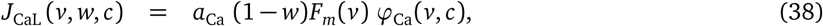

where *c* represents the intracellular Ca^2+^ concentration, and φ_*x*_ is a difference of exponential functions as defined above, with *x* ∈ {NaK, NaCa, KD, CaL}. The temporal evolution for *v* is then described by

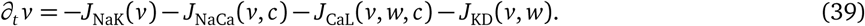

During pacemaking the concentrations of Na^+^ and K^+^ across the membrane are assumed to change negligibly, but the Ca^2+^ concentration does change at least 10-fold (Rasmusson et al., 1990a,b; Herrera-Valdez and Lega, 2011). Therefore, the system includes an equation for the dynamics for *c* in which *c* converges to a steady state *c*_*∞*_ in the absence of Ca^2+^ fluxes, and increases proportionally to the total transport of Ca^2+^ ions via L-type channels and Na^+^-Ca^2+^ exchangers (Fig. 4). Explicitly,

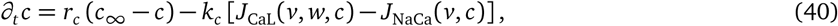

where *k*_*c*_ (*µ*M/mV) represents the impact of the transmembrane Ca^2+^ fluxes on the free intracellular Ca^2+^ concentration. The minus sign in front of *k*_*c*_ accounts for the fact that the sign of the *J*_CaL_ is negative. The sign in front of *J*_NaCa_ is because the forward flux of Ca^2+^ mediated by the Na-Ca exchanger is opposite to that of electrodiffusive Ca^2+^.

The solutions of equations (32)-(40) with parameters as in Table 2 reproduce important features of the membrane dynamics observed in the rabbit’s central sinoatrial node, including the period (*ca*. 400 ms), amplitude (*ca*. 70 mV), and maximum *∂*_*t*_ *v* (<10 V/s) of the action potentials (Zhang et al., 2000).

**Table 2.** **Parameters for the cardiac SAN pacemaker model. The amplitudes** *a*_*l*_ **can be thought of as a 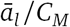 where** *C*_*M*_ **is a constant that represents the rate of change in charge around the membrane as a function of** *v***, and** *l* ∈ {Ca*L*, K, NaK, NaCa}.

| Parameter | Value | Units | Description |
| --- | --- | --- | --- |
| $C_M$ | 30 | pF | Membrane capacitance |
| $\bar{a}_{\text{Ca}}$ | 1 | pA | Maximum amplitude for the L-type $\text{Ca}^{2+}$ current |
| $\bar{a}_{\text{K}}$ | 700 | pA | Maximum amplitude for the $\text{K}^+$ current |
| $\bar{a}_{\text{NaK}}$ | 1 | pA | Maximum amplitude for the $\text{Na}^+$-$\text{K}^+$ current |
| $\bar{a}_{\text{NaCa}}$ | 2.5 | pA | Maximum amplitude for the $\text{Na}^+$-$\text{Ca}^{2+}$ current |
| $a_{\text{Ca}} = \bar{a}_{\text{Ca}}/C_M$ | 0.0333 | pA/pF | Maximum amplitude for the L-type $\text{Ca}^{2+}$ current |
| $\bar{a}_{\text{K}} = \bar{a}_{\text{K}}/C_M$ | 23.3333 | pA/pF | Maximum amplitude for the $\text{K}^+$ current |
| $\bar{a}_{\text{NaK}} = \bar{a}_{\text{NaK}}/C_M$ | 0.03333 | pA/pF | Maximum amplitude for the $\text{Na}^+$-$\text{K}^+$ current |
| $\bar{a}_{\text{NaCa}} = \bar{a}_{\text{NaCa}}/C_M$ | 0.1 | pA/pF | Maximum amplitude for the $\text{Na}^+$-$\text{Ca}^{2+}$ current |
| $v_{\text{ATP}}$ | -420 | mV | Potential ATP hydrolysis |
| $v_{\text{Na}}$ | 60 | mV | Nernst potential for $\text{Na}^+$ |
| $v_{\text{K}}$ | -89 | mV | Nernst potential for $\text{K}^+$ |
| $v_{\text{NaK}} = 3v_{\text{Na}} - 2v_{\text{K}} + v_{\text{ATP}}$ | -62 | mV | Reversal potential for the for $\text{Na}^+$-$\text{K}^+$ ATPase current |
| $v_{\text{NaCa}} = 2v_{\text{Ca}} - 3v_{\text{Na}}$ | - | mV | Reversal potential for the for $\text{Na}^+$-$\text{Ca}^{2+}$ current ($v_{\text{Ca}}$ depends continuously on $[\text{Ca}]_i$) |
| $v_{m13}$ | -18 | mV | Half-activation potential for $\text{Ca}_{v13}$ L-type $\text{Ca}^{2+}$-current |
| $v_w$ | 0 | mV | Half-activation potential for the transient $\text{K}^+$-current |
| $g_{m13}$ | 4 | - | Activation slope factor for the $\text{Ca}_{v13}$ L-type $\text{Ca}^{2+}$-current |
| $g_w$ | 3 | - | Activation slope factor for the $\text{K}^+$-current |
| $r_w$ | 0.05 | $\text{s}^{-1}$ | Activation rate for the cardiocyte $\text{K}^+$-current |
| $b_w$ | 0.35 | - | Activation slope factor for the $\text{K}^+$-current |
| $b_{\text{NaK}}$ | 0.5 | - | Non-rectification bias for the $\text{Na}^+$-$\text{K}^+$-current |
| $b_{\text{K}}$ | 0.5 | - | Rectification for the transient $\text{K}^+$-current |
| $b_{\text{Na}}$ | 0.5 | - | Non-rectification bias for the transient $\text{Na}^+$-current |
| $b_{\text{Ca}}$ | 0.5 | - | Non-rectification bias for the $\text{Ca}_{v13}$ L-type $\text{Ca}^{2+}$-current |
| $c_{\infty}$ | 0.1 | $\mu\text{M}$ | Minimal (resting) intracellular $\text{Ca}^{2+}$-concentration |
| $r_c$ | 0.02 | $\text{ms}^{-1}$ | $\text{Ca}^{2+}$ removal rate |
| $k_c$ | 0.00554 | - | Conversion factor between $\text{Ca}^{2+}$ current and intracellular $\text{Ca}^{2+}$ concentration |

The solutions of the system show a number of interesting features related to ionic fluxes. First, the Na-Ca current reverses when *v* = *v*_NaCa_ (Fig. 4A, blue line). During the initial depolarization and until the maximum downstroke rate, approximately, *v*_NaCa_ *< v*, which means *J*_NaCa_ > 0, so that Ca^2+^ extrusion by the Na-Ca exchanger occurs only for a brief period of time during the downstroke and also after each action potential (Fig. 4C, blue line). Second, as previously reported in different studies involving spiking dynamics, the time course of the Ca^2+^ current shows a partial inactivation with a double peak (Fig. 4B, blue line) around a local minimum (Rasmusson et al., 1990a,b; Carter and Bean, 2009), and in agreement with data from voltage-clamp experiments (Mangoni et al., 2006). A number of models have made attempts to reproduce the double activation by making extra assumptions about gating (Rasmusson et al., 1990a,b). For instance, some models include a second activation variable, or the multiple terms in the steady state gating, or in the time constant for activation or inactivation. However, the explanation for the double peak can be much simpler. The calcium current *J*_CaL_ is a negative-valued, non monotonic function for *v < v*_Ca_, which can be thought of as a product of a amplitude term that includes gating and the function *ϕ*_CaL_. The normalized current *J*_CaL_ has a local minimum (maximum current amplitude) around −10 mV (Fig. 4B, blue line and Fig. 5A, blue line), after which the current decreases, reaching a local maximum as the total current passes through zero, at the peak of the action potential around 10 mV, (Fig. 4B, where *∂*_*t*_ *v* = 0). The first peak for the Ca^2+^ current occurs when *v* reaches the maximum depolarization rate (Fig. 5B). As *v* increases (e.g. upstroke of the action potential). The second peak for the current occurs as the membrane potential decreases, and passes again through the region where the maximal current occurs (local minimum for *J*_*Ca*_). The two local minima for *J*_CaL_ represent peaks in the Ca^2+^ current that have different amplitudes due the difference in the time course of *v* during the upstroke and the downstroke of the action potential (Fig. 5A, blue line, and Fig. 5B, where *∂*_*t*_ *v* = 0). It is important to remark that the dual role played by *w* is not the cause of the double activation. This is illustrated by analyzing the behavior of a non-inactivating *J*_CaL_ without the inactivation component, (Fig. 5A, gray line). The double activation can also be observed in models in which the activation of K^+^ channels and the inactivation of Ca^2+^ or Na^+^ channels are represented by different variables (Rasmusson et al., 1990a) and in dynamic voltage clamp experiments on neurons in which there are transient and persistent sodium channels (Carter and Bean, 2009).

**Figure 5.**
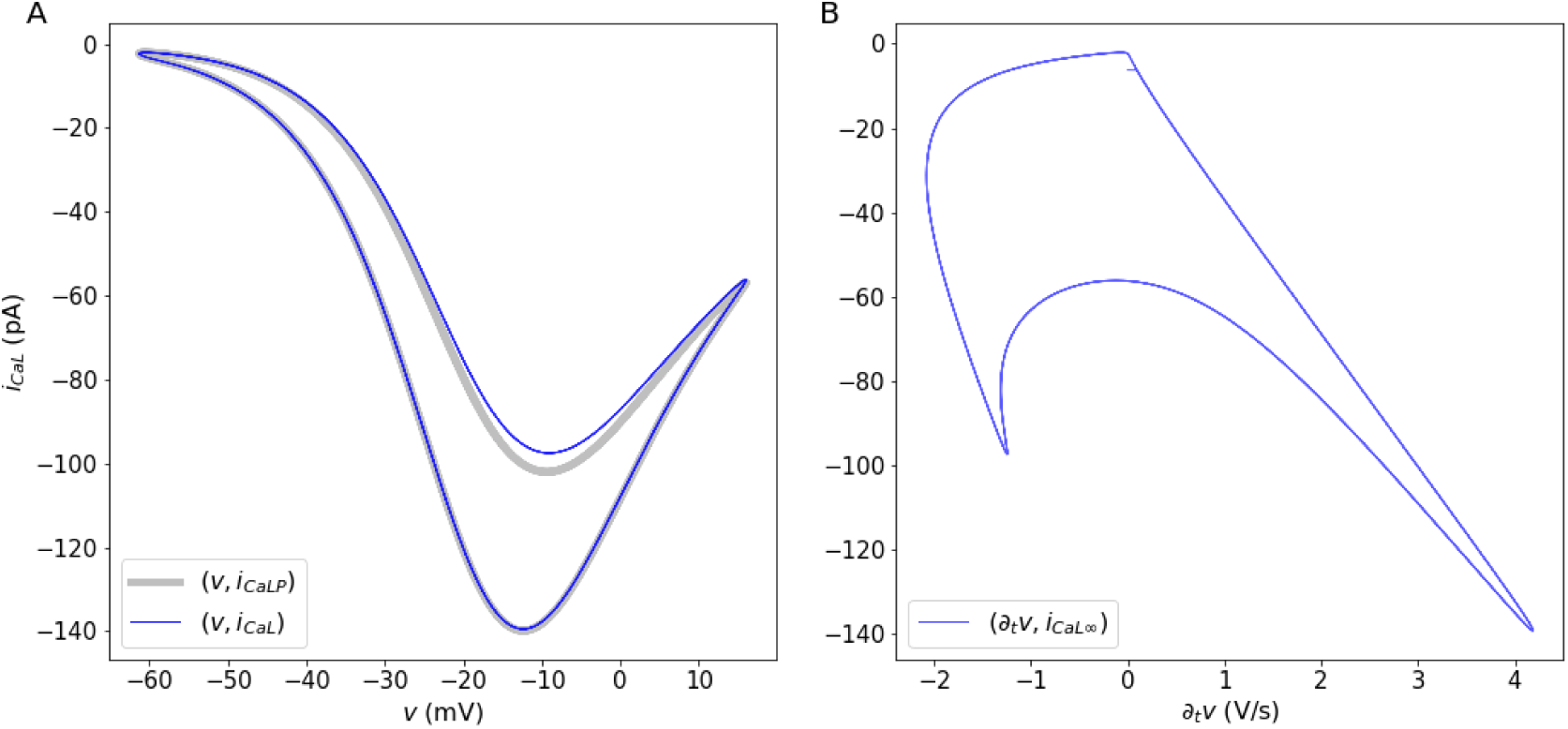
**Dynamics of the calcium current and double activation. A. Behavior of the inactivating L-type Ca**^2+^ **current with respect to the transmembrane potential (blue line) and a non-inactivating current (gray line), and (B) with respect to the time-dependent change in** *v*.

The double peak in the Ca^2+^ current is reflected in the intracellular Ca^2+^ concentration (Figure 6, gray line), and by extension, on the Nernst potential for Ca^2+^ (Figure 6, blue line), which display two increasing phases and two decreasing phases, respectively. The first and faster phase in both cases occur during the initial activation of the L-type channels. The second phase occurs during the downstroke, as second peak of the Ca^2+^ current occurs. As a consequence, the reversal potential for the Na-Ca exchanger, *v*_NaCa_ = 3*v*_Na_ *-*2*v*_Ca_ (Figure 6, orange line) also has two phases, this time increasing. Increasing the intracellular Ca^2+^ (Figure 6, gray line) concentration decreases the Nernst potential for Ca^2+^, and viceversa. By extension, the reversal potential for the Na-Ca exchanger, *v*_NaCa_ = 3*v*_Na_ *-*2*v*_Ca_ becomes larger when *c* increases. Ca^2+^ enters the cell in exchange for Na^+^ that moves out when *v > v*_NaCa_, during most of the increasing phase and the initial depolarization phase of the action potential (blue lines in Figure 4A and C, and Figure 6).

**Figure 6.**
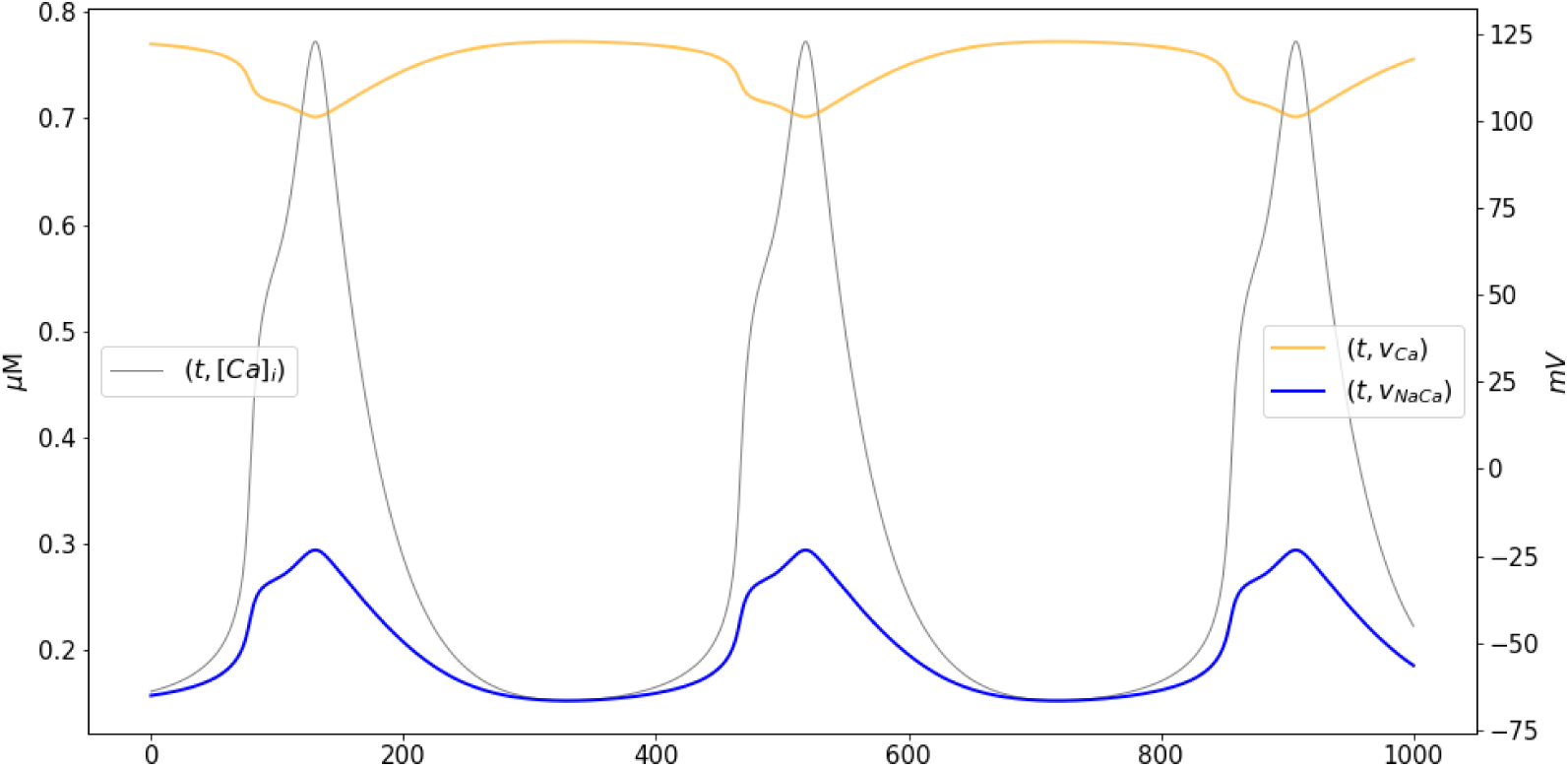
**Calcium dynamics during pacemaking. Time courses of the intracellular calcium concentration (gray, left axis), the Nernst potential for Ca**^2+^ **(orange, right axis), and the reversal potential for the Na-Ca exchanger (blue, right axis)**.

## Discussion

A generic, macroscopic model for transmembrane fluxes has been derived by directly calculating the work required to transport molecules across the membrane. The derivation is based on a general thermodynamic scheme that takes into account the rate, stoichiometry, and the direction in which the molecules are transported across the membrane. These biophysical parameters are then combined to write expressions for directional fluxes based on van’t Hoff (1884) and Arrhenius (1889) formulations, weighted as in the Butler/Erdey-Gruz/Volmer equation (Butler, 1924; Erdey-Grúz and Volmer, 1930). The result is a general description (equation 16) of the transmembrane molecular flux as a difference of exponential functions, that describes the transport dynamics in the “forward” and “backward” directions, relative to a source compartment. The two exponential functions depend on a common expression involving the transmembrane concentrations of the molecules being transported, and possibly the transmembrane potential when transport is electrogenic.

Rectification, an asymmetry in the flow, is typically modeled modifying the dynamics of the gating variables of the current. The general formulas for transmembrane transport include a bias term *b* that controls the relative contribution of inward and outward components the transport. Hence, different types of rectification can be described by favoring one of the directions for transport, conceptually in line with the “anomalous rectification” originally reported by Katz (1949) for K^+^ in muscle cells. The bias term is not part of any gating mechanism. Instead, it represents the asymmetry in bidirectional flux Based on the work of Riedelsberger et al. (2015), the inward (outward, respectively) rectification in K^+^ channels occurs when the fourth transmembrane segment of the channel (S4) is located closer to the intracellular (extracellular) portion of the membrane in its open configuration. There are other reports that show that asymmetries in bidirectional transport occur as a consequence of changes in the three dimensional structure of the protein mediating the transport (Halliday and Resnick, 1981; Quistgaard et al., 2013). Therefore, the rectification term can be thought of as representing a structural component of the transmembrane protein through which molecules move (Fig. 2). Outward rectification in K^+^ channels can be explained, for instance, by biasing the flux of K^+^ the forward (outward) direction (*b*_K_ > 1/2). Instead, inward rectification can be obtained by biasing the transport in the backward (inward) direction (*b*_K_ < 1/2). It is important to remark that non-rectifying currents with *b* = 1/2 are nonlinear functions of Δ*G*, which shows that the nonlinearity of the current-voltage relationships is not the defining characteristic of rectification; as argued in some textbooks (see Kew and Davies, 2010).

The formulation for transmembrane flux may be rewritten in different alternative forms that can be found throughout the literature (see equations (13) and (14), Goldman, 1943; Johnston et al., 1995). Of particular interest, the widely used conductance-based models for current from the seminal work of Hodgkin and Huxley (1952) turn out to be linear approximations of the generic current described here (Herrera-Valdez, 2012, 2014). This explains why the Hodgkin and Huxley (1952) model is captures many of the defining features of action potential generation, in spite of modeling ionic currents as resistive. Another interesting case is that electrodiffusive transmembrane currents derived from the Nernst-Planck equation (Nernst, 1888; Planck, 1890), turn out particular cases of the generic formulation presented here (see also Herrera-Valdez, 2014, for details). Examples include the constant field approximation (Hille, 1992; Johnston et al., 1995; Clay et al., 2008), the non-rectifying currents proposed by Endresen et al. (2000), and more general electrodiffusive currents that includes a bias term accounting for rectification (Johnston et al., 1995; Herrera-Valdez, 2014).

Possibly of interest to mathematicians working on bifurcation theory, a third order approximation (equation (17)) resembling the Fitz-Hugh equations (FitzHugh, 1955, 1961; Fitz-Hugh, 1966), can be used to construct models that give very close approximations to the full model, while keeping biophysical characteristics like rectification and the multiplicative interaction between the slow variable *w* and the fast variable *v*. Further, the third order approximation opens the possibility of expanding on the analysis of dynamical systems based on these generic formulas to study normal forms and bifurcations. Another possible use of the third order approximations is in the construction of network models.

One question of interest because of its possible impact on the interpretation of results from existing modeling studies is how does the excitability and the resulting dynamics in a model of membrane dynamics change when using the thermodynamic transmembrane currents or their approximations? The question has been addressed in a study in which two simple neuronal models with currents mediated by Na^+^ and K^+^, each equipped with the same biophysical gating properties and the same relative contributions for the currents, but one with currents as in equation (19), the other with conductance-based currents. The two models display a number of qualitative and quantitative differences worth considering while making the choice of a model in theoretical studies (Herrera-Valdez, 2012). For a start, the two models are not topologically equivalent across many ratios of the relative contributions of K^+^ and Na^+^ channels (Herrera-Valdez, 2012); as would be expected by the fact that conductance-based formulations are only linear approximations of the generic currents. One of the most notable differences is the contribution of the nonlinear, high order terms from equation (16), which results in more realistic upstrokes for action potentials and an overall increased excitability; in this case characterized in terms of the minimum sustained current necessary to produce at least one action potential. The increased excitability of the membrane is due, in part, to the large, exponential contribution of the open Na^+^ and Ca^2+^ channels, but not the K^+^ channels, to the change in the transmembrane potential near rest. The time course of the Na^+^ current during the beginning of the action potential with the generic model is much sharper than that of the conductance-based formulation, resulting in a faster upstroke of the action potential; and in better agreement with observations in cortex and other tissues (Naundorf et al., 2006). It is important to remark that the sharper increase in the change of the membrane potential is a consequence of the nonlinear driving force terms of the current (the flux term in the generic formulation) and not in the activation dynamics for the transient Na^+^ current.

The generic formulation for both passive and active transmembrane transport can be thought of as a tool that facilitates the construction and analysis of models of membrane potential dynamics. The generality and versatility of the thermodynamic transmembrane transport formulations is illustrated with a model of the dynamics of cardiac pacemaking (equations (39)-(34)). Another example with a model for a fast spiking interneuron can be found in the Appendix. The ion fluxes in the model are assumed to be mediated by two different types of voltage-gated channels and two different types of pumps, all represented with the *same* functional form (see Herrera-Valdez and Lega (2010); DiFrancesco and Noble (1985); Rasmusson et al. (1990b) for examples in which that is not the case).

One important advantage of the generic formulation is that it includes the possibility of explicitly estimating the number of channels or pumps mediating each of the transport mechanisms of interest. This has proven to be useful to study the relative contributions of different currents to the excitability of neurons (see Herrera-Valdez et al., 2013), cardiocytes (Herrera-Valdez, 2014), and other different tissues (unpublished work). Another extension of possible interest is that of modelling the transmembrane transport between organelles and the cytosolic compartment, which can be done by directly replacing the difference *c*_*s*_*- d*_*s*_ in equation (1) with 1 or −1, accounting for the direction of transmembrane motion of molecules relative to the outer compartment. This and other generalizations enable the possibility of studying the interdependence between electrical excitability across tissues and animal species (Herrera-Valdez et al., 2013), and its cross-interactions with metabolism and other processes of physiological importance, all from a general theoretical framework with common formulations.

### Implications for experimentalists

One of the main advantages of the generic expressions is that fits to ionic currents can be made straight from the voltage-clamp data without much effort, and without having to calculate conductances, which amounts to imposing the assumption that the current to voltage relationship is linear. Fits to experimental currents can then be directly put into equations describing the change in the membrane potential, and model membrane dynamics of interest without having to make many extra adjustments, as it is the case for most conductance-based models restricted to data.

The model for current in equation (19) has been used to construct simplified models for the membrane dynamics of different cell types using experimental data. Examples include motor neurons in Drosophila melanogaster (Herrera-Valdez et al., 2013), pyramidal cells in the young and ageing hippocampus of rats (McKiernan et al., 2015), medium spiny neurons in the mouse striatum (Suárez et al., 2015), rabbit sinoatrial node cells (Herrera-Valdez, 2014), and other types of excitable cells (McKiernan and Herrera-Valdez, 2012).

## Conclusions

A generic model that describes physiological transmembrane transport of molecules has been derived by considering basic thermodynamical principles. The model unifies descriptions of transport mediated by channels and pumps, it can model biases in either one of the directions of flow, and it can be easily converted into a model for current in the case of electrogenic transport. As it is desirable in all models, the generic expressions can be thought of as extensions of some previous models. In particular, it is shown that the conductance-based model for current turns out to be a first order approximation of the generic formulation.

The expressions for current and molecular fluxes across the membrane based on the generic formulation can be used to build general models of transmembrane potential using a unified framework (Shou et al., 2015).

### Author contributions

MAHV had the original idea, developed and analyzed the models, performed the numerical experiments, and wrote the article.

### Grant information

This work was supported by UNAM-PAPIIT IA208618.

## Acknowledgements

The author wishes to thank Joceline Lega, Timothy Secomb, and Raphael Gruener at the University of Arizona; Jose Bargas-Diaz and Antonio Laville from the Cellular Physiology Institute at UNAM; and Erin C. McKiernan from the Physics Department at UNAM for all the time spent in discussions that helped to solidify and deepen the ideas presented in this paper.

---

**Table A1.** **Physical constants. The conversion factor** f **from pA to** *µM ·ms*^*-*1^ = *mM ·s*^*-*1^ **implies** ⇒ *M* = *f ·*10^*-*9^**Coul. Then** *f* = 10^9^ **M/Coul =** 10^9^ *·* 96485.3329*/F* ≈ 10^14^ **M/Coul**.

| Constant | Value | Units | Description |
| --- | --- | --- | --- |
| $T_0$ | 273.16 | degrees Kelvin | Zero absolute temperature |
| $N_A$ | $6.023 \times 10^{23}$ | molecules/Mole | Avogadro's number |
| $q$ | $1.60217733 \times 10^{-19}$ | Coulombs/molecule | Elementary charge |
| $k$ | $1.381 \times 10^{-23}$ | Joules/$^{\circ}$Kelvin | Boltzmann constant |
| $F = qN_A$ | 96485.33289 | Coulombs/Mole | Faraday's constant (the magnitude of electric charge per mole of electrons) |
| $R = kN_A$ | 1.987 | cal/(Mole $^{\circ}$Kelvin) | Gas constant |

## Appendix

### The Goldman constant field approximation from the general formulation

Let *x* = [*x*]_*j*_, *j ∈* {0, 1} (extra, and intracellular concentrations of an ion of type *x*. The Goldman-Hodgkin-Katz equation describing the transmembrane current carried by *x*-ions is given by

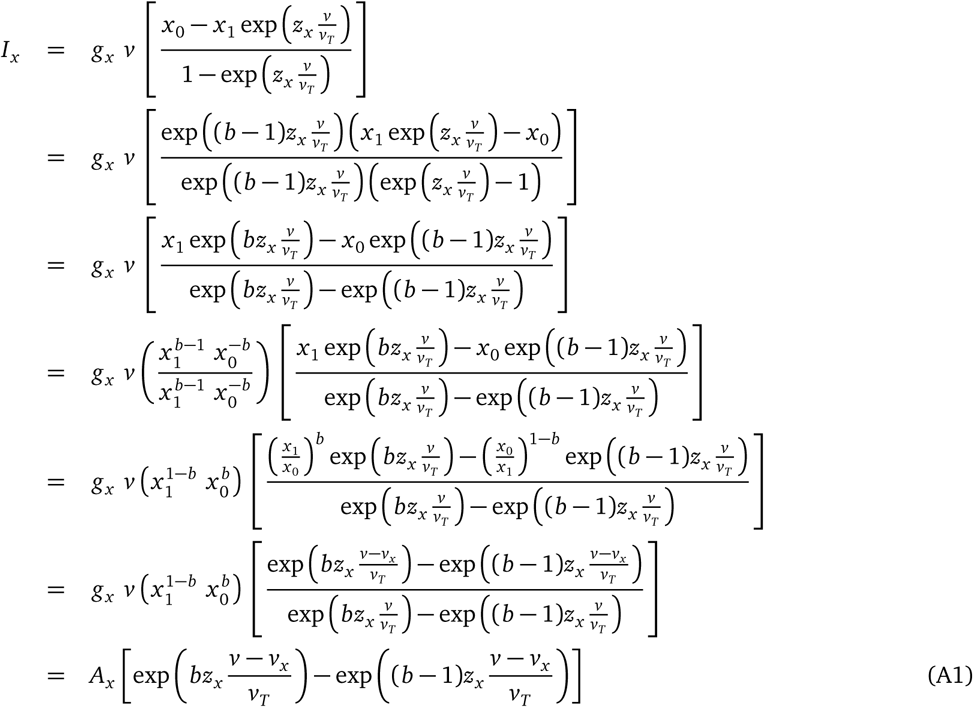

where

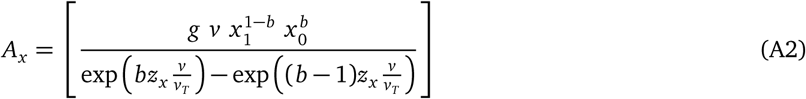

is an amplitude term that can be approximated by a constant (Nonner et al., 1998; Nonner and Eisenberg, 1998; Endresen et al., 2000). A specific example for a calcium current at 24°C can be found in equations 25 and 26 in the article by Herrera-Valdez and Lega (2011). Notice that equations (A1)-(A2) reduce to

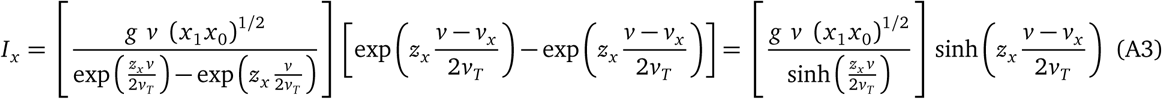

when *b* = 1/2.

### Fast spiking interneuron dynamics

A simple model of the dynamics of a fast spiking (FS) striatal interneuron can be constructed using (31). To do so, assume the transmembrane potential depends on three currents respectively mediated by Na-K pumps, non-inactivating K^+^ channels, and Na^+^ channels with transient dynamics, with voltage-dependent gating in both channels. It is also assumed that the proportion of activated K^+^ is represented by a variable *w* ∈ [0, 1], which also represents the proportion of inactivated Na^+^ channels (Av-Ron et al., 1991; Rinzel, 1985). That is, 1 *-w* represents the proportion of non-inactivated Na^+^ channels. The dynamics for *w* can be assumed to follow a logistic scheme, capturing the behaviour of delayed-rectifier K^+^ currents typically recorded in voltage clamp mode without adding extra powers to *w* (see for instance Hodgkin and Huxley, 1952, and the Appendix). It is also assumed that sodium channel activation is fast, described by its steady state function of *v*.

**Figure A1.**
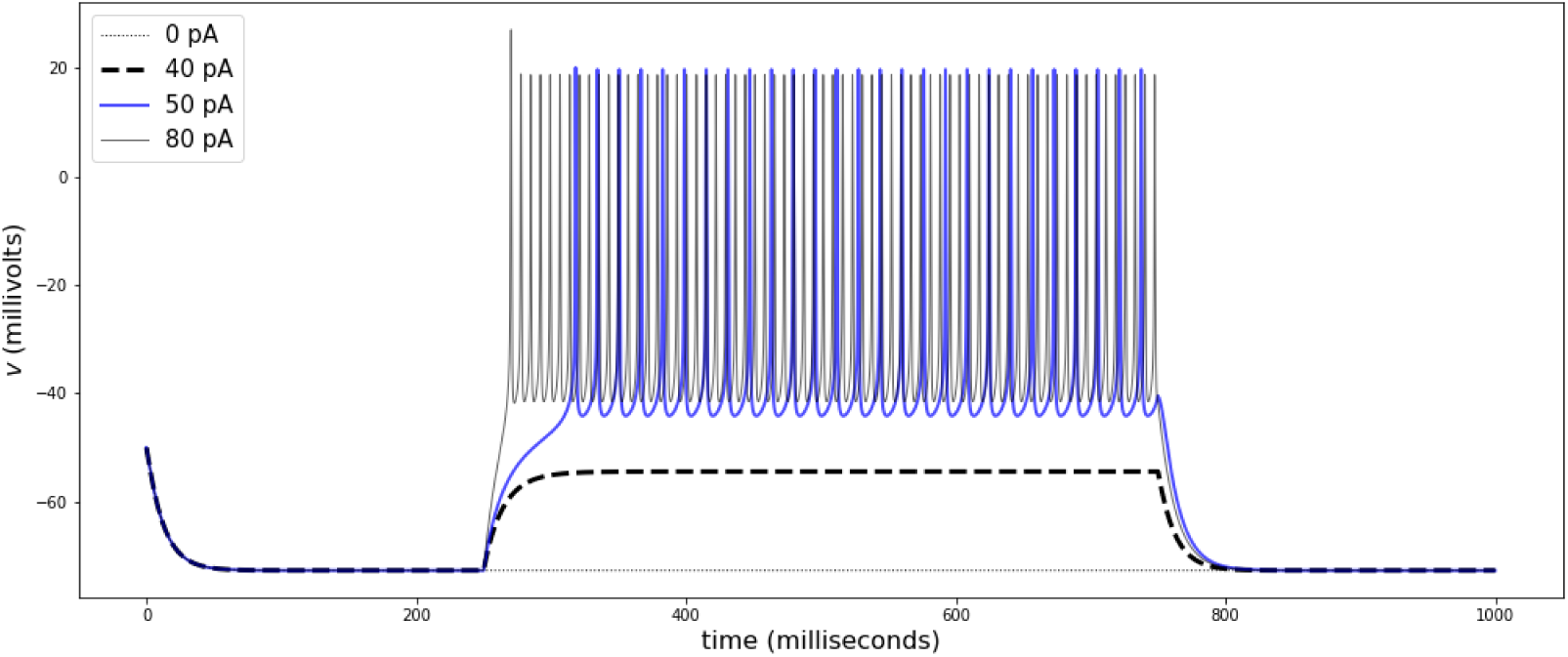
Rest to spiking transitions of FS interneuron under current clamp. The traces show responses to current-clamp stimulation of different amplitudes. The transition between rest and spiking with a rheobase occurs between 40 and 50 pA, as shown for some FS neurons in the mouse striatum (Orduz et al., 2013). The traces correspond to stimulation amplitudes of 0 (gray dots), 40 (black dashed line), 50 (blue), and 80 pA (gray). Parameters can be found in Table A2.

Explicitly, the dynamics of the system can then be captured by coupled differential equations of the form

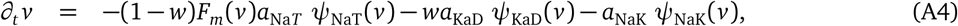

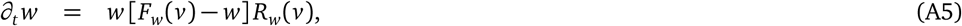

The activation rate for K^+^ channels depends is a voltage-dependent functions *R*_*w*_ and *F*_*w*_ as defined for the cardiac pacemaking model.

The dynamics of the system are such that, as *v* increases, *w* increases, but at a slower rate in comparison to *v*. This is because the activation *w* is always moving toward its steady state value, which increases as *v* increases. Once *w* increases, the Na^+^ current tends to decrease and the K^+^ current tends to increase, thereby causing a decrease in *v*. The slower dynamics in *w* relative to those in *v* capture the delay between the amplification caused by the Na^+^ current and the recovery caused by the K^+^ current. The current mediated by Na/K-ATPase acts as an extra attracting force toward *v*_NaK_ that increases nonlinearly as the distance between *v* and *v*_NaK_ increases. Striatal FS interneurons display maximum *∂*_*t*_ *v* between 100 and 200 V/s. In current clamp mode, most neurons are silent, and show transitions between rest and repetitive spiking at a rheobase current of approximately 90 pA, with initial firing rates between 50 and 60 Hz and a delay to first spike in the transition that decreases as the stimulus amplitude increases (Fig.A1, parameters in Table A2).

To include these properties into the model, the membrane capacitance was specified first, then the maximum *∂*_*t*_ *v* was adjusted by fitting the parameter *a*_NaT_, and then the contributions for the K^+^ channels and the Na-K ATPase are set to obtain spiking and fit the rheobase.

The model in equations (A4)-(A5) reproduces dynamics observable in fast spiking neurons in CA1 (Erisir et al., 1999) or in the striatum (Orduz et al., 2013; Tepper et al., 2010).

**Table A2.** Parameters for the fast spiking interneuron model.

| Parameter | Value | Units | Description |
| --- | --- | --- | --- |
| Current amplitudes and capacitance for the neuronal membrane model |  |  |  |
| $C_m$ | 30 | pF | Membrane capacitance |
| $\bar{a}_{NaK}$ | 67 | pA | Maximum amplitude for the $Na^+$-$K^+$ ATPase current |
| $\bar{a}_K$ | 4400 | pA | Maximum amplitude for the delayed-rectifier $K^+$ current |
| $\bar{a}_{Na}$ | 1400 | pA | Maximum amplitude for the transient $Na^+$ current |
| $v_{ATP}$ | -430 | mV | Potential ATP hydrolysis |
| $v_{NaK} = 3v_{Na} - 2v_K + v_{ATP}$ | -72 | mV | Reversal potential for the for $Na^+$-$K^+$ ATPase current |
| $v_K$ | -89 | mV | Nernst potential for $K^+$ |
| $v_{Na}$ | 60 | mV | Nernst potential for $Na^+$ |
| $v_{mT}$ | -17 | mV | Half-activation potential for the transient $Na^+$-current |
| $v_w$ | -5 | mV | Half-activation potential for the transient $K^+$-current |
| $g_{mT}$ | 5 | – | Activation slope factor for the transient $Na^+$-current |
| $g_w$ | 4 | – | Activation slope factor for the $K^+$-current |
| $r_w$ | 2 | $s^{-1}$ | Activation rate for the neuronal $K^+$-current |
| $b_w$ | 0.3 | – | Activation slope factor for the $K^+$-current |
| $b_{NaK}$ | 0.5 | – | Non-rectification for the $Na^+$-$K^+$-current |
| $b_K$ | 0.5 | – | Non-rectification for the transient $K^+$-current |
| $b_{Na}$ | 0.5 | – | Non-rectification for the transient $Na^+$-current |

## Footnotes

1 The transmembrane potential for which there is a zero net flux of s-ions across the membrane, as given by the Nernst-Planck equation, is 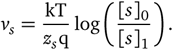

## References

Wilfred D Stein and Thomas Litman. Channels, carriers, and pumps: an introduction to membrane transport. Elsevier, 2014.

H Stanley Bennett. The concepts of membrane flow and membrane vesiculation as mechanisms for active transport and ion pumping. The Journal of biophysical and biochemical cytology, 2(4):99, 1956.

Andreas Blicher and Thomas Heimburg. Voltage-gated lipid ion channels. PLoS One, 8(6):e65707, 2013.

B Hille. Ionic Channels of Excitable Membranes. Sinauer Associates, Sinauer Associates, Inc. Sunderland, Mass. 01375, 1992.

Isabelle Favre, Edward Moczydlowski, and Laurent Schild. On the structural basis for ionic selectivity among na+, k+, and ca2+ in the voltage-gated sodium channel. Biophysical journal, 71(6):3110–3125, 1996.

W. Almers and E. W. McCleskey. Non-selective conductance in calcium channels of frog muscle: calcium selectivity in a single-file pore. The Journal of Physiology, 353(1):585, 1984.

Declan A Doyle, Joao Morais Cabral, Richard A Pfuetzner, Anling Kuo, Jacqueline M Gulbis, Steven L Cohen, Brian T Chait, and Roderick MacKinnon. The structure of the potassium channel: molecular basis of k+ conduction and selectivity. science, 280(5360):69–77, 1998.

Hans H Ussing. The active ion transport through the isolated frog skin in the light of tracer studies. Acta Physiologica, 17(1): 1–37, 1949a.

Hans H Ussing. Transport of ions across cellular membranes. Physiological reviews, 29(2):127–155, 1949b.

J Chr SKou. Enzymatic basis for active transport of na+ and k+ across cell membrane. Physiological Reviews, 45(3):596–618, 1965.

Hans H Ussing. The distinction by means of tracers between active transport and diffusion. Acta Physiologica, 19(1):43–56, 1949c.

D.C. Gadsby. Ion channels versus ion pumps: the principal difference, in principle. Nature Reviews Molecular Cell Biology, 10 (5):344–352, 2009.

Marco Arieli Herrera-Valdez and Joceline Lega. Reduced models for the pacemaker dynamics of cardiac cells. Journal of Theoretical Biology, 270(1):164–176, 2010. doi: 10.1016/j.jtbi.2010.09.042.

D. E. Goldman. Potential, impedance, and rectification in membranes. Journal of general Physiology, 27(1):37, 1943.

L. Barr. Membrane potential profiles and the Goldman equation. Journal of Theoretical Biology, 9(3):351–356, 1965.

K. S. Cole. Electrodiffusion models for the membrane of squid giant axon. Physiological Reviews, 45(2):340, 1965. ISSN 0031-9333.

Douglas B Kell. On the functional proton current pathway of electron transport phosphorylation: an electrodic view. Biochimica et Biophysica Acta (BBA)-Reviews on Bioenergetics, 549(1):55–99, 1979.

P Läuger. Ion transport through pores: a rate-theory analysis. Biochimica et Biophysica Acta (BBA)-Biomembranes, 311(3): 423–441, 1973.

C. F. Stevens and R. W. Tsien. Ion permeation through membrane channels, volume 3. Raven Press, 1979.

Philippa M Wiggins. The relationship between pump and leak: Part 1. application of the butler-volmer equation. Bioelectrochemistry and Bioenergetics, 14(4):313–326, 1985a.

Philippa M Wiggins. Relationship between pump and leak: Part 2. a model of the na, k-atpase functioning both as pump and leak. Bioelectrochemistry and Bioenergetics, 14(4):327–337, 1985b.

Philippa M Wiggins. Relationship between pump and leak: Part 3. electrical coupling of na+-solute uptake to the na, k-atpase. Bioelectrochemistry and Bioenergetics, 14(4-6):339–345, 1985c.

D. DiFrancesco and D. Noble. A model of cardiac electrical activity incorporating ionic pumps and concentration changes. Philosophical Transactions of the Royal Society of London. Series B, Biological Sciences, 307:353–398, 1985.

L. P. Endresen, K. Hall, J. S. Hoye, and J. Myrheim. A theory for the membrane potential of living cells. European Journal of Biophysics, 29:90–103, 2000.

Eduardo Marbán. Cardiac channelopathies. Nature, 415(6868):213, 2002.

Frances M Ashcroft. Atp-sensitive potassium channelopathies: focus on insulin secretion. The Journal of clinical investigation, 115(8):2047–2058, 2005.

A. L. Hodgkin and A. F. Huxley. A quantitative description of membrane current and its application to conduction and excitation in nerve. Journal of Physiology, 117:500–544, 1952.

Alan L Hodgkin and Bernard Katz. The effect of sodium ions on the electrical activity of the giant axon of the squid. The Journal of physiology, 108(1):37–77, 1949.

William F Pickard. Generalizations of the goldman-hodgkin-katz equation. Mathematical biosciences, 30(1-2):99–111, 1976.

Th Rosenberg and W Wilbrandt. The kinetics of membrane transports involving chemical reactions. Experimental cell research, 9(1):49–67, 1955.

R. L. Rasmusson, J. W. Clark, W. R. Giles, K Robinson, R. B. Clark, E. F. Shibata, and D. L. Campbell. A mathematical model of electrophysiological activity in the bullfrog atrial cell. Am. J. Physiol., 259:H370–H389, 1990a.

R. L. Rasmusson, J. W. Clark, W. R. Giles, E. F. Shibata, and D. L. Campbell. A mathematical model of bullfrog cardiac pacemaker cell. Am. J. Physiol., 259:H352–H369, 1990b.

William F Pickard. A postulational approach to the problem of ion flux through membranes. Mathematical Biosciences, 4(1-2): 7–21, 1969.

John A Jacquez and Stanley G Schultz. A general relation between membrane potential, ion activities, and pump fluxes for symmetric cells in a steady state. Mathematical Biosciences, 20(1-2):19–25, 1974.

John A Jacquez. A general relation between membrane potential, ion activities, and pump fluxes for nonsymmetric cells in a steady state. Mathematical Biosciences, 53(1-2):53–57, 1981.

D. Johnston, S. M. S. Wu, and R. Gray. Foundations of cellular neurophysiology. MIT press Cambridge, MA, 1995. ISBN 0262100533.

Marco Arieli Herrera-Valdez. Geometry and nonlinear dynamics underlying electrophysiological phenotypes in biophysical models of membrane potential. Dissertation. Ph.D. in Mathematics. University of Arizona, 2014.

P De Weer, David C Gadsby, and RF Rakowski. Voltage dependence of the na-k pump. Annual Review of Physiology, 50(1): 225–241, 1988.

David J. Aidley. The Physiology of Excitable Cells. Cambridge University Press, 4 edition, 1998. ISBN 0521574153,9780521574150. URL http://gen.lib.rus.ec/book/index.php?md5=25AD083C33F37AC8E44F3CE90E3B3B84.

M.P. Blaustein, J.P.Y. Kao, and D.R. Matteson. Cellular physiology. Elsevier/Mosby, 2004. ISBN 0323013414.

Walther Nernst. Zur kinetik der in lösung befindlichen körper. Zeitschrift für physikalische Chemie, 2(1):613–637, 1888.

C Tanford. Equilibrium state of atp-driven ion pumps in relation to physiological ion concentration gradients. The Journal of general physiology, 77(2):223–229, 1981.

LJ Mullins. The generation of electric currents in cardiac fibers by na/ca exchange. American Journal of Physiology-Cell Physiology, 236(3):C103–C110, 1979.

LA Venetucci, AW Trafford, SC O’neill, and DA Eisner. Na/ca exchange. Annals of the New York Academy of Sciences, 1099(1): 315–325, 2007.

AK Sen and WF Widdas. Determination of the temperature and ph dependence of glucose transfer across the human erythrocyte membrane measured by glucose exit. The Journal of physiology, 160(3):392–403, 1962.

Dorine M Starace, Enrico Stefani, and Francisco Bezanilla. Voltage-dependent proton transport by the voltage sensor of the shakerk+ channel. neuron, 19(6):1319–1327, 1997.

David T Yue, Peter H Backx, and John P Imredy. Calcium-sensitive inactivation in the gating of single calcium channels. Science, 250(4988):1735–1738, 1990.

Béla Novák and John J Tyson. Design principles of biochemical oscillators. Nature reviews Molecular cell biology, 9(12):981, 2008.

JAV Butler. Studies in heterogeneous equilibria. part 2.—the kinetic interpretation of the nernst theory of electromotive force. Transactions of the Faraday Society, 19(March):729–733, 1924.

Th Erdey-Grúz and M Volmer. Zur theorie der wasserstoff überspannung. Zeitschrift für Physikalische Chemie, 150(1):203–213, 1930.

Richard Courant and Fritz John. Introduction to calculus and analysis I. Springer Science & Business Media, 2012.

Michael Spivak. Calculus on manifolds: a modern approach to classical theorems of advanced calculus. CRC Press, 2018.

B Katz. Les constantes electriques de la membrane du muscle. Arch Sci Physiol, 3:285–299, 1949.

Clay M Armstrong and Leonard Binstock. Anomalous rectification in the squid giant axon injected with tetraethylammonium chloride. The Journal of general physiology, 48(5):859–872, 1965.

RH Adrian. Rectification in muscle membrane. Progress in biophysics and molecular biology, 19:341–369, 1969.

JW Woodbury. Eyring rate theory model of the current-voltage relationships of ion channels in excitable membranes. Advances in Chemical Physics: Chemical Dynamics: Papers in Honor of Henry Eyring, Volume 21, pages 601–617, 1971.

Janin Riedelsberger, Ingo Dreyer, and Wendy Gonzalez. Outward rectification of voltage-gated k+ channels evolved at least twice in life history. PloS one, 10(9):e0137600, 2015.

Michael Hollmann, Melissa Hartley, and Stephen Heinemann. Ca2+ permeability of ka-ampa–gated glutamate receptor channels depends on subunit composition. Science, 252(5007):851–853, 1991.

Allan G Lowe and Adrian R Walmsley. The kinetics of glucose transport in human red blood cells. Biochimica et Biophysica Acta (BBA)-Biomembranes, 857(2):146–154, 1986.

J. D. Hunter. Matplotlib: A 2d graphics environment. Computing In Science & Engineering, 9(3):90–95, 2007. doi: 10.1109/MCSE.2007.55.

Robert L Post and Philip C Jolly. The linkage of sodium, potassium, and ammonium active transport across the human erythrocyte membrane. Biochimica et biophysica acta, 25:118–128, 1957.

PJ Garrahan and IM Glynn. The behaviour of the sodium pump in red cells in the absence of external potassium. The Journal of physiology, 192(1):159–174, 1967.

David C Gadsby, Junko Kimura, and Akinori Noma. Voltage dependence of na/k pump current in isolated heart cells. Nature, 315(6014):63–65, 1985.

J Brian Chapman. On the reversibility of the sodium pump in dialyzed squid axons: A method for determining the free energy of atp breakdown? The Journal of general physiology, 62(5):643, 1973.

Masakazu Nakao and David C Gadsby. [na] and [k] dependence of the na/k pump current-voltage relationship in guinea pig ventricular myocytes. The Journal of General Physiology, 94(3):539–565, 1989.

Kanako Hamada, Hiroshi Matsuura, Mitsuru Sanada, Futoshi Toyoda, Mariko Omatsu-Kanbe, Atsunori Kashiwagi, and Hitoshi Yasuda. Properties of the na+/k+ pump current in small neurons from adult rat dorsal root ganglia. British journal of pharmacology, 138(8):1517–1527, 2003.

Sanda Despa, Mohammed A Islam, Christopher R Weber, Steven M Pogwizd, and Donald M Bers. Intracellular na+ concentration is elevated in heart failure but na/k pump function is unchanged. Circulation, 105(21):2543–2548, 2002.

BJ Pitts. Stoichiometry of sodium-calcium exchange in cardiac sarcolemmal vesicles. coupling to the sodium pump. Journal of Biological Chemistry, 254(14):6232–6235, 1979.

John P Reeves and Calvin C Hale. The stoichiometry of the cardiac sodium-calcium exchange system. Journal of Biological Chemistry, 259(12):7733–7739, 1984.

Wolfgang Nonner and Robert Eisenberg. Ion permeation and glutamate residues linked by poisson-nernst-planck theory in l-type calcium channels. Biophysical Journal, 75:1287–1305, 1998.

Matteo E Mangoni, Brigitte Couette, Emmanuel Bourinet, Josef Platzer, Daniel Reimer, Jörg Striessnig, and Joël Nargeot. Functional role of l-type cav1. 3 ca2+ channels in cardiac pacemaker activity. Proceedings of the National Academy of Sciences, 100(9):5543–5548, 2003.

L. Sanders, S. Rakovic, M. Lowe, P. A. D. Mattick, and D. A. Terrar. Fundamental importance of Na+–Ca2+ exchange for the pacemaking mechanism in guinea-pig sino-atrial node. The Journal of physiology, 571(3):639, 2006.

T. Shibasaki. Conductance and kinetics of delayed rectifier potassium channels in nodal cells of the rabbit heart. The Journal of Physiology, 387(1):227, 1987.

E. Av-Ron, H. Parnas, and L. A. Segel. A minimal biophysical model for an excitable and oscillatory neuron. Biological Cybernetics, 65(6):487–500, 1991.

S. Tsunoda and L. Salkoff. The major delayed rectifier in both Drosophila neurons and muscle is encoded by Shab. Journal of Neuroscience, 15(7):5209–5221, 1995.

M Covarrubias, A Wei, and L Salkoff. Shaker, Shal, Shab, and Shaw expresss independent K-current systems. Neuron(Cambridge, Mass.), 7(5):763–773, 1991.

A. R. Willms, D. J. Baro, R. M. Harris-Warrick, and J. Guckenheimer. An improved parameter estimation method for hodgkinhuxley models. Journal of Computational Neuroscience, 6(2):145–168, 1999.

Marco Arieli Herrera-Valdez and Joceline Lega. Reduced models for the pacemaker dynamics of cardiac cells. Journal of Theoretical Biology, 270(1):164–176, 2011.

H Zhang, AV Holden, I Kodama, H Honjo, M Lei, T Varghese, and MR Boyett. Mathematical models of action potentials in the periphery and center of the rabbit sinoatrial node. American Journal of Physiology-Heart and Circulatory Physiology, 279(1): H397–H421, 2000.

Brett C Carter and Bruce P Bean. Sodium entry during action potentials of mammalian neurons: incomplete inactivation and reduced metabolic efficiency in fast-spiking neurons. Neuron, 64(6):898–909, 2009.

Matteo E Mangoni, Brigitte Couette, Laurine Marger, Emmanuel Bourinet, Jörg Striessnig, and Joël Nargeot. Voltage-dependent calcium channels and cardiac pacemaker activity: from ionic currents to genes. Progress in biophysics and molecular biology, 90(1):38–63, 2006.

Jacobus Henricus van’t Hoff. Etudes de dynamique chimique, volume 1. Muller, 1884.

Svante Arrhenius. Über die reaktionsgeschwindigkeit bei der inversion von rohrzucker durch säuren. Zeitschrift für physikalische Chemie, 4(1):226–248, 1889.

David Halliday and Robert Resnick. Fundamentals of physics. John Wiley & Sons, 1981.

Esben M Quistgaard, Christian Löw, Per Moberg, Lionel Trésaugues, and Pär Nordlund. Structural basis for substrate transport in the glut-homology family of monosaccharide transporters. Nature structural & molecular biology, 20(6):766, 2013.

James NC Kew and Ceri H Davies. Ion channels: from structure to function. Oxford University Press, USA, 2010.

Marco Arieli Herrera-Valdez. Membranes with the same ion channel populations but different excitabilities. PloS one, 7(4): e34636, 2012.

Max Planck. Ueber die potentialdifferenz zwischen zwei verdünnten lösungen binärer electrolyte. Annalen der Physik, 276(8): 561–576, 1890.

J. R. Clay, D. Paydarfar, and D. B. Forger. A simple modification of the hodgkin and huxley equations explains type 3 excitability in squid giant axons. Journal of The Royal Society Interface, 5(29):1421–1428, 2008.

Richard FitzHugh. Mathematical models of threshold phenomena in the nerve membrane. Bulletin of Mathematical Biology, 17(4):257–278, 1955.

Richard FitzHugh. Impulses and physiological states in theoretical models of nerve membrane. Biophysical journal, 1(6): 445–466, 1961.

Richard Fitz-Hugh. Mathematical models of excitation and propagation in nerve. Publisher Unknown, 1966.

Björn Naundorf, Fred Wolf, and Maxim Volgushev. Unique features of action potential initiation in cortical neurons. Nature, 440(7087):1060, 2006.

Marco Arieli Herrera-Valdez, Erin Christy McKiernan, Sandra Daniela Berger, Stephanie Ryglewski, Carsten Duch, and Sharon Crook. Relating ion channel expression, bifurcation structure, and diverse firing patterns in a model of an identified motor neuron. Journal of Computational Neuroscience, pages 1–19, 2013.

Erin Christy McKiernan, Marco Arieli Herrera-Valdez, and Diano Fabio Marrone. A biophysical, minimal model to explore age-related changes in ion channel gene expression and excitability in ca1 pyramidal cells. Society for Neurosciences Annual Meeting, Session 628: Learning and Memory: Aging III, Poster 628.10/AA45., 2015.

Paola Suárez, Marco Arieli Herrera-Valdez, José Bargas, and Elvira Galarraga. Un modelo biofísico de neuronas estriatales de proyección que toma en cuenta la contribución de canales de calcio cav3. Escuela de Otoño de Biomatemáticas, Jalapa, Veracruz, México., 2015.

E.C. McKiernan and Marco Arieli Herrera-Valdez. From spinal cord to hippocampus: links between bifurcation structure, ion channel expressio n, and firing patterns in a variety of neuron types. BMC Neuroscience, 13(Suppl 1):P121, 2012.

Wenying Shou, Carl T Bergstrom, Arup K Chakraborty, and Frances K Skinner. Theory, models and biology. Elife, 4:e07158, 2015.

W. Nonner, D. P. Chen, and Robert Eisenberg. Anomalous mole fraction effect, electrostatics, and binding in ionic channels. Biophysical journal, 74(5):2327–2334, 1998. ISSN 0006-3495.

J. Rinzel. Excitation dynamics: insights from simplified membrane models. In Fed. Proc. 44, volume 2944, 1985.

David Orduz, Don Patrick Bischop, Beat Schwaller, Serge N Schiffmann, and David Gall. Parvalbumin tunes spike-timing and efferent short-term plasticity in striatal fast spiking interneurons. The Journal of physiology, 591(13):3215–3232, 2013.

A Erisir, D Lau, B Rudy, and CS Leonard. Function of specific k+ channels in sustained high-frequency firing of fast-spiking neocortical interneurons. Journal of neurophysiology, 82(5):2476–2489, 1999.

James M Tepper, Fatuel Tecuapetla, Tibor Koós, and Osvaldo Ibáñez-Sandoval. Heterogeneity and diversity of striatal gabaergic interneurons. Frontiers in neuroanatomy, 4:150, 2010.

